# Visualizing conformational dynamics of proteins in solution and at the cell membrane

**DOI:** 10.1101/294538

**Authors:** Sharona E. Gordon, Mika Munari, William N. Zagotta

## Abstract

Conformational dynamics underlie enzyme function, yet are generally inaccessible via traditional structural approaches. FRET has the potential to measure conformational dynamics *in vitro* and in intact cells, but technical barriers have thus far limited its accuracy, particularly in membrane proteins. Here, we combine amber codon suppression to introduce a donor fluorescent noncanonical amino acid with a new, biocompatible approach for labeling proteins with acceptor transition metals in a method called ACCuRET (Anap Cyclen-Cu^2+^ resonance energy transfer). We show that ACCuRET measures absolute distances and distance changes with high precision and accuracy using maltose binding protein as a benchmark. Using cell unroofing, we show that ACCuRET can accurately measure rearrangements of proteins in native membranes. Finally, we implement a computational method for correcting the measured distances for the distance distributions observed in proteins. ACCuRET thus provides a flexible, powerful method for measuring conformational dynamics in both soluble proteins and membrane proteins.

## Introduction

Structural dynamics of proteins, particularly those at cell membranes, underlie many cell-signaling events (1). Structural rearrangements in membrane proteins can occur in response to binding of extracellular or intracellular ligands, covalent modification (e.g. phosphorylation), changes in membrane voltage, and mechanical forces in the membrane. These rearrangements, in turn, regulate enzyme activity, open pores, and transport molecules across the membrane. In this way, membrane proteins serve as molecular transducers that mediate the communication and transport between the cell and the external world. The molecular mechanisms, however, for these transduction processes remain largely unknown.

The number of solved membrane protein structures has grown exponentially in the last 30 years, and there are now more than 750 unique structures in the protein structure database (http://blanco.biomol.uci.edu/mpstruc/; accessed 3/29/18). These structures provide high-resolution information on the position of each atom in the protein, but provide little information about the functional states, number of states, relative energies of the states, or allowed transitions. In contrast, functional measurements like electrophysiology provide high-resolution information on states and energetics, but little information on structure. In addition, electrophysiology gives an excellent readout on rearrangements in the pore domain but is a poor surrogate for probing structural and energetic rearrangements in other regions, e.g. agonist/antagonist binding sites. This state of affairs has left a gap in our ability to link structure and function, as we often don’t know the functional state for a particular structure or the structural state(s) that underlie a particular function.

To fill this gap a method is needed that allows us to measure the structural dynamics and energetics of membrane proteins, preferably in their native membrane environment. Many of the existing methods for measuring structural dynamics have limitations. Recent advances in cryo-electron microscopy have greatly facilitated the measurement of multiple protein conformations, often in the same sample (2). However, these conformational states can only be distinguished if they have substantial structural differences, and their functional state and energetics are still unknown. Magnetic resonance methods have proven useful for measuring structural dynamics but require high concentrations of purified protein, a particular challenge for membrane proteins. Finally, fluorescence resonance energy transfer (FRET) provides a sparse measurement of protein structure and can be recorded on a biological time scale (milliseconds to seconds) on small quantities of protein (even single molecules) in their native environment (3, 4). Standard FRET, however, suffers from nonspecific labeling, inappropriate distance range for intramolecular measurements, and inaccurate measurements of distances and distance changes (5, 6).

Our goal was to establish and optimize a method for measuring and analyzing the structural dynamics of membrane proteins in their native membrane environment. Our approach, which we call ACCuRET (<u>A</u>nap <u>C</u>yclen <u>Cu</u>^2+^ <u>R</u>esonance <u>E</u>nergy <u>T</u>ransfer), combines together three technologies: 1) transition metal ion FRET (tmFRET) for accurately measuring interatomic distances, 2) amber codon suppression to specifically label the protein with a fluorescent, noncanonical amino acid as the FRET donor, and 3) a novel orthogonal and biocompatible method for introducing a specific, a high-affinity binding site for a metal ion that serves as the FRET acceptor.

To measure interatomic distances we have used tmFRET between a fluorophore and a transition metal divalent cation (7-9). Transition metal cations, such as Ni^2+^, Co^2+^, and Cu^2+^, act as non-fluorescent FRET acceptors, quenching the fluorescence of donor fluorophores with an efficiency that is highly distance dependent (10-14). Thus, tmFRET provides a sparse, dynamic measurement of structure on minute amounts of functioning proteins in their native environment. tmFRET has a number of significant advantages over standard FRET methods for measuring rearrangements in proteins (3). 1) tmFRET has a working range of about 10-20 Å, much shorter than standard FRET, and on the order of the distance between proximal structural elements in proteins. 2) tmFRET is steeply distance dependent, but has little orientation dependence because the metal ion has multiple transition dipole moments (15). 3) The method utilizes minimal metal binding sites, making the position of the metal a more faithful representation of the position of the protein backbone. 4) Metal binding is rapid and reversible so the fraction of fluorescence quenched at a saturating concentration of metal gives the absolute FRET efficiency and therefore the distance. And 5) metals with different R_0_ values (distance producing 50% FRET efficiency) can be chosen to tune the measurement to the distance of interest.

The power of using fluorescence to probe protein dynamics within the cell is limited by the specificity with which a protein can be labeled with a fluorophore. For measuring the structure and dynamics of a protein, it is also important to use a small fluorophore attached to the protein with a short linker (3). To achieve this specific fluorescent labeling with a small fluorophore, we used amber codon suppression to introduce the fluorescent, noncanonical amino acid L-Anap (16). We chose L-Anap, a derivative of prodan, because of its small size (about the same as the amino acid tryptophan) and useful spectral properties (16, 17). By introducing a fluorophore as the side-chain of an amino acid, we eliminated the need for a linker whose distance from the protein backbone and flexibility reduces the accuracy and empirical sensitivity of FRET (5, 6, 9, 18). L-Anap is therefore useful as a tmFRET donor for distance measurements.

Membrane proteins are fully functional only in their native membrane environment. For many membrane proteins, studying the relevant structural rearrangements demands access to the cytoplasmic side of the protein for introduction of ligands, probes, etc. To study membrane proteins, we have recently implemented cell unroofing, an approach borrowed from the EM literature (19), as a medium-throughput platform for measuring tmFRET from membrane proteins in native cell membranes (17, 20). This technique utilizes a mechanical sheering force to dislodge the dorsal surface of cells and remove all soluble cellular contents and organelles, leaving the ventral surface of the cells intact as plasma membrane sheets containing their native lipids and membrane proteins. Cell unroofing provides access to the intracellular surface of the bilayer for application of transition metals and intracellular ligands in an environment more physiological than membrane proteins reconstituted into synthetic liposomes and much higher throughput than inside-out patch-clamp recording.

Previous studies using FRET on membrane proteins have produced uncertain results about the size of structural rearrangements, often underestimating changes in distance (3). Here we validate our tmFRET approach using maltose binding protein (MBP) as a model system. MBP is a bacterial periplasmic protein that undergoes a significant structural rearrangement upon binding maltose (Fig. 1A; Supplemental Movie 1). The structure and rearrangement of MBP has been extremely well characterized, with over 260 structures in the protein structure database to date, both with and without ligands bound. Using both soluble and membrane-bound forms of MBP, we have determined the precision and accuracy of ACCuRET for measuring intramolecular distances and changes in distance due to ligand binding (Fig. 1, Supplemental Movie 1). Using MBP we show that: 1) ACCuRET provides an accurate measurement of intramolecular distances between 10 and 20 Å; 2) ACCuRET measures small changes in distance; 3) L-Anap can be introduced efficiently and specifically into proteins; 4) minimal metal binding sites can be introduced using a small, reversible cysteine reactive metal chelate; and 5) similar ACCuRET measurements can be made in membrane proteins in unroofed cells. These experiments establish this approach as a powerful method to measure the structural dynamics of membrane proteins.

**Fig. 1.**
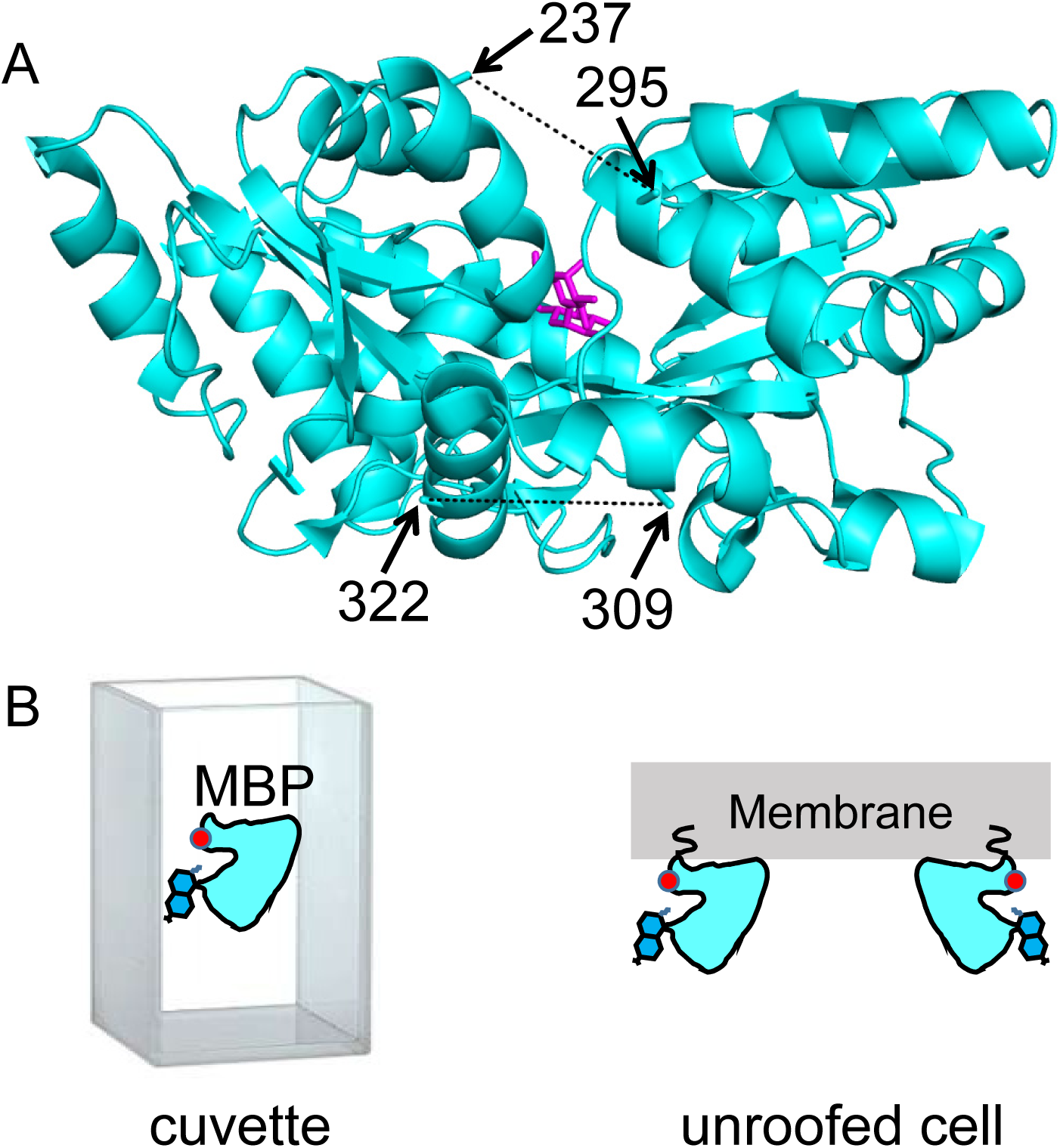
MBP as a model system for optimizing and validating ACCuRET. (A) X-ray crystal structure of MBP bound to maltose (PDB ID: 1N3X) showing the positions of the donor and acceptor FRET pairs at the top (295 and 237, respectively) and bottom (322 and 309, respectively) of the clamshell, with the distance represented by dashed lines. The distance separating the 295/237 FRET pair decreases and the 322/309 FRET pair increases with maltose binding (Supplemental Movie 1). (B) Soluble MBP was studied in a cuvette in a fluorometer (left), and membrane-bound MBP was studied in membranes of unroofed cells (right).The red dot represents bound metal, the blue side-chain represents Anap, and the squiggle at the membrane represents protein lipidation at the engineered CAAX site.

## Results

### Selection of FRET pairs on Maltose Binding Protein

MBP is a clamshell shaped protein that undergoes a significant closure of the clamshell upon binding its ligand, maltose (Fig. 1A; Supplemental Movie 1). In addition, maltose binding leads to a smaller opening of the backside of the clamshell. By introducing one FRET pair at the top of the clamshell and a second FRET pair on the backside of the clamshell, we can address the accuracy and precision of ACCuRET to measure absolute distances as well as ligand-dependent increases and decreases in distance.

Our criteria for selection of FRET pairs were 1) the sites are solvent exposed, 2) the sites are on rigid secondary structural elements (α helices in MBP), 3) the distance between the sites is predicted to fall in the working range of tmFRET (~10-20 Å), and 4) the distance between sites undergoes a moderate change between apo and holo MBP. Our first FRET pair was at positions 295 (donor) and 237 (acceptor) on the outer lip of the clamshell (Fig. 1A; Supplemental Movie 1). From β carbon distances in the X-ray structures, these sites are 21 Å apart in the apo state and 13 Å apart in the holo state, a distance change of about −8 Å (shortening) with maltose binding. Our second FRET pair was at positions 322 (donor) and 309 (acceptor) on the backside of the clamshell (Fig. 1A; Supplemental Movie 1). These sites are 13 Å apart in the apo state and 17 Å apart in the holo state, a distance change of +4 Å (lengthening) with maltose binding. In this study, we used these two FRET pairs to evaluate the accuracy and precision of ACCuRET with different acceptor metal-binding sites and different metals as well as for MBP in solution and bound to the membrane.

### Specific labeling with fluorophore and transition metal ion

ACCuRET requires the protein to be site-specifically labeled with a donor fluorophore and an acceptor transition metal ion. To achieve specific labeling of MBP with a small donor fluorophore, we used amber codon suppression to introduce the fluorescent noncanonical amino acid L-Anap (Fig. 2A) (16, 21). Our amber codon suppression method required the use of two plasmids (17, 22). One plasmid encodes MBP with a TAG stop codon (the amber codon) engineered at the site for L-Anap incorporation. The second plasmid, pAnap, developed by Peter Schultz’s lab (16), encodes an evolved tRNA and amino acyl tRNA synthetase, which the cell uses to produce L-Anap-loaded tRNA complementary to the TAG codon (Fig. 2A). HEK293T/17 cells were co-transfected with the two plasmids and incubated in a cell-permeable methyl ester (ME) version of L-Anap.

**Fig. 2.**
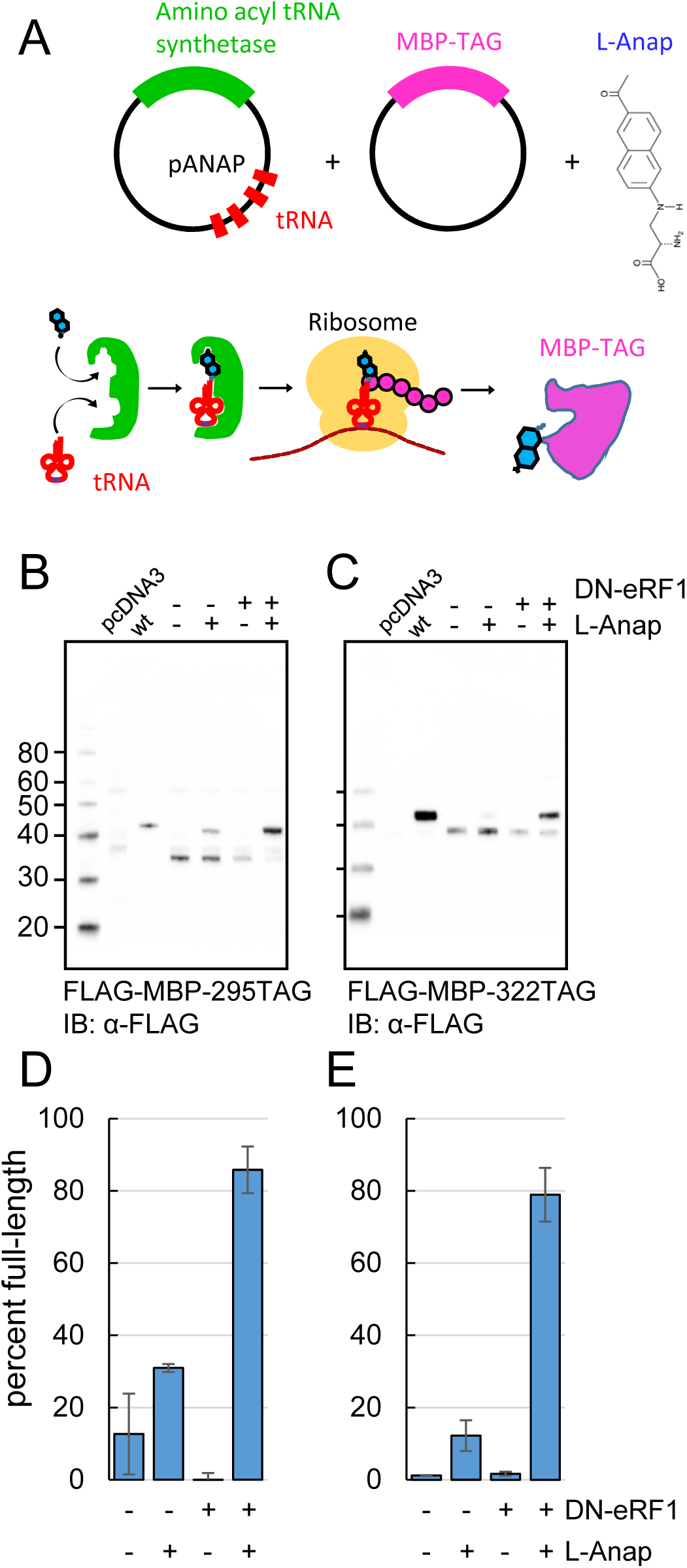
Incorporation of the fluorescent noncanonical amino acid Anap into MBP as the FRET donor. (A) Cartoon representation of amber codon suppression strategy for site-specific incorporation of L-ANAP into MBP. (B-C) Full-length protein for MBP-TAG constructs, where TAG indicates that the construct includes an internal amber stop codon, was produced with L-ANAP-ME and increased with DN-eRF1 cotransfection. Western blots are shown of clarified lysates of cells cotransfected with (B) MBP-295TAG or (C) MBP-322TAG, pANAP, and either with (+) or without (-) DN-eRF1, and cultured with (+) or without (−) L-ANAP-ME in the medium. The blots were probed with an anti-FLAG antibody that recognizes both truncated and full length MBP. Vector only transfected cells (pcDNA3) were used as a negative control and wild-type MBP (wt) was used as a positive control. The amount of wild-type MBP loaded on the gel was less than the amount of the TAG mutations. As discussed in Materials and Methods, all MBP constructs used in this work included an N-terminal FLAG epitope. (D-E) Quantitation of the percent of full-length product of (D) MBP-295TAG or (E) MBP-322TAG under the same conditions as B and C. Shown are mean ± SEM with n = 3.

We achieved specific incorporation of L-Anap at donor positions 295 and 322 (Fig. 1A). MBP-Anap was the primary band visible in gels of crude cell lysates using in-gel L-Anap fluorescence (Supplementary Fig. 1). In addition, Western blots of FLAG-MBP containing the K295TAG or E322TAG mutation revealed full-length MBP only when the cells were incubated with L-Anap-ME (Fig. 2B, C). However, even when L-Anap was present in the medium, a moderate amount of truncated product, in which translation terminated at the introduced stop codon, was observed for both MBP-295TAG and MBP-322TAG, as shown by the plots in Fig. 2D, E (fraction of full-length MBP was 30% and 10% respectively). These results indicate that L-Anap incorporation at positions 295 and 322 in MBP in HEK293T/17 cells was specific, but not particularly efficient.

To increase the efficiency of L-Anap incorporation, we co-expressed the MBP-TAG and pAnap plasmids with a third plasmid encoding a dominant negative form of eukaryotic Release Factor 1 (DN-eRF1). Previously, Jason Chin’s group showed that DN-eRF1 increases the efficiency of incorporation of noncanonical amino acids in mammalian cells (23), presumably by delaying release of the translation product from the ribosome when encountering an amber stop codon. For both sites in MBP, coexpression with DN-eRF1 substantially increased the fraction of full-length product from cells incubated with L-Anap-ME (from 30 to 86% for MBP-295TAG and from 10 to 79% for MBP-322TAG) (Fig. 2D, E). Coexpression with DN-eRF1 also increased the absolute amount of full-length protein relative to that observed with wild-type MBP, from 1.6% (±0.4%; n=3) to 6.3% (±0.5%; n=3) for MBP-295TAG and from 5.5% (±4%; n=3) to 50% (±27%; n=3) for MBP-322TAG. We observed little or no read-through (i.e. incorporation of a natural amino acid at the stop codon), even when co-expressing DN-eRF1, as little full length MBP-TAG protein was observed when L-Anap was omitted from the cell growth medium (Fig. 2B, C). These results establish a method for achieving specific, high-efficiency fluorescent labeling of MBP for ACCuRET experiments.

L-Anap’s small size, short linker, and spectral properties makes it useful for tmFRET experiments (17, 22, 24, 25). The peak absorption wavelength of L-Anap is 360 nm (16). The emission of L-Anap is environmentally sensitive, blue shifting as the environment becomes less polar. The emission peak of L-Anap was 494 nm in our buffer and 475 nm in ethanol (see Materials and Methods, Fig. 13A, B). For the MBP-295Anap and MBP-322Anap in this study, the peak emissions were 491 nm and 478 nm, respectively, suggesting that neither 295Anap nor 322Anap is fully solvent exposed and the that environment of 322Anap is more hydrophobic than that of 295Anap. We found that the quantum yield of L-Anap was also site dependent, measured to be 0.31 for MBP-295Anap and 0.47 for MBP-322Anap (see Materials and Methods and Fig. 13). L-Anap’s low quantum yield, together with its low extinction coefficient (~20×10^3^ M^−1^cm^−1^; (16, 17)), makes L-Anap relatively dim compared to most visible light fluorophores; however, its fluorescence intensity is sufficient for most macroscopic fluorescence experiments. L-Anap’s emission spectrum overlaps the absorption spectra of many transition metal ions (such as Ni^2+^, Co^2+^, and Cu^2+^) making it a useful FRET donor for tmFRET (Fig. 3C).

**Fig. 3.**
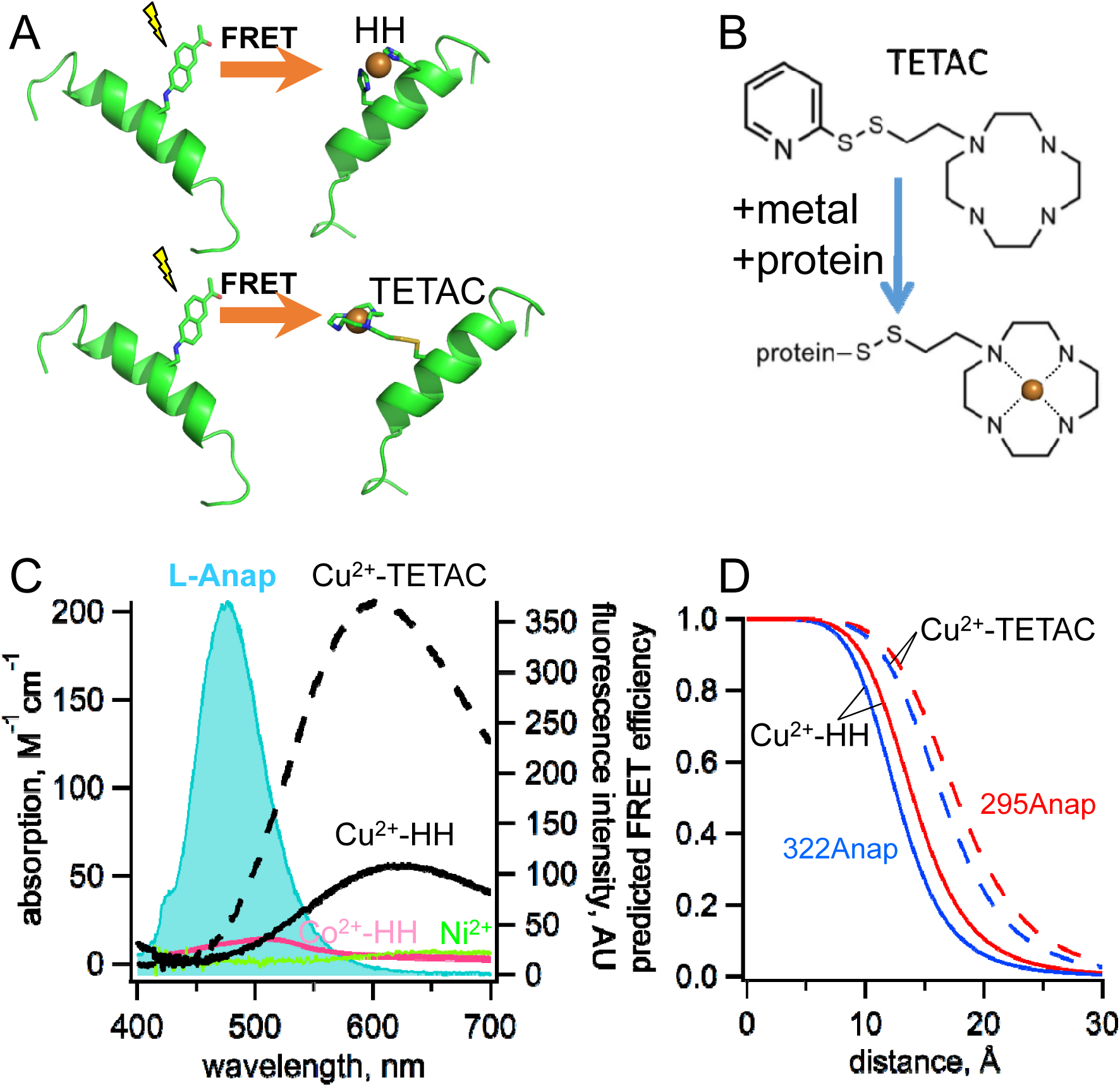
Incorporation of transition metal ions as the FRET acceptor using standard HH sites and TETAC. (A) Diagram illustrating tmFRET between Anap (left) and transition metal (right) for two different types of metal binding sites: a standard HH site (top) and TETAC site (bottom). (B) Structure of TETAC, binding of Cu^2+^ (orange ball), and reaction with a cysteine in a protein. (C) Spectral properties of free L-ANAP and Cu^2+^-TETAC make them an ideal FRET pair for measuring small distances on the order of 10–20 Å. The emission spectrum from L-Anap (magenta) is overlaid on the absorption spectra of Cu^2+^-TETAC (dashed black line), Cu^2+^-HH (black line), Co^2+^-HH (magenta line) and Ni^2+^ (green line). (D) Distance dependence of the FRET efficiency predicted with the Förster equation for each of the two FRET pairs, MBP-295Anap (red line) and MBP-322Anap (blue line), for both Cu^2+^-TETAC (dashed line) and Cu^2+^-HH (solid line).

The standard method to introduce a transition metal ion acceptor for tmFRET uses a dihistidine (HH) motif introduced at positions *i* and *i*+4 on an α helix (Fig. 3A) or at i and *i*+2 on a beta strand (8, 9, 12, 26-28). These minimal metal binding sites can bind Ni^2+^, Co^2+^, and Cu^2+^ with affinities in the 1 to 100 µM range, and binding can be reversed with EDTA. Although this method places the transition metal ion very close to the protein backbone, the metal concentrations required are high, so that they are not biocompatible and are subject to off-target binding.

To circumvent the limitations of standard HH sites, we developed a new, better method to introduce a specific, high affinity metal-binding site in a protein by labeling a cysteine with 1-(2-pyridin-2-yldisulfanyl)ethyl)-1,4,7,10-tetraazacyclododecane (TETAC; Fig. 3A,B). TETAC is a cysteine-reactive compound with a short linker to a cyclen ring that binds transition metal ions with very high-affinity (sub-nanomolar). Single cysteines can then be introduced in the protein at sites of interest and reacted with TETAC to create a mixed disulfide linkage to a cyclen metal-binding site.

### ACCuRET measures FRET and ligand-dependent changes in FRET

ACCuRET uses TETAC to introduce minimal metal binding sites. TETAC has many advantages compared with standard HH sites for tmFRET: 1) it requires only nominal concentrations of free Cu^2+^ eliminating off-target binding of Cu^2+^ to endogenous binding sites on the protein; 2) it can be used with any type of secondary structural element; 3) its reaction with endogenous cysteines produces tmFRET only for sites that are very close (< 20 Å), so endogenous cysteines need not be eliminated; 4) it can be easily removed with reducing agent, making its use compatible with physiological concentrations of other metals, e.g. Ca^2+^; and 5) it significantly blue shifts and increases the absorbance of Cu^2+^, extending the range of R_0_ values available for tmFRET (Fig. 3C). As shown in Fig. 3D, the distance dependence of the FRET efficiency predicted by the Förster equation for each of our two L-Anap sites in MBP is shifter by almost 4 Å to longer distances for Cu^2+^-TETAC compared to Cu^2+^-HH. ACCuRET therefore offers a flexible and biocompatible method for measuring distances in the biological distance range.

For each of our two FRET pairs (Fig. 1A), we made three constructs: one with no metal binding site, one with a standard HH binding site, and one with a Cys, for modification by TETAC. For MBP-295Anap, the HH was introduced at positions 233 and 237 (MBP-295Anap-HH) and the Cys was introduced at position 237 (MBP-295Anap-C). For MBP-322Anap, the HH was introduced at positions 305 and 309 (MBP-322Anap-HH) and the Cys was introduced at position 309 (MBP-322Anap-C). For both donor sites, control constructs were also made with no introduced metal binding sites (MBP-295Anap and MBP-322Anap) to measure other forms of energy transfer and nonspecific decreases in fluorescence, and correct the FRET efficiency measurements accordingly (see Materials and Methods). These constructs were all expressed in HEK293T/17 cells as amino-terminal FLAG fusions and purified using anti-FLAG beads for subsequent analysis (Supplemental Fig. 1).

ACCuRET was readily observed in both MBP-Anap-C constructs. We recorded emission spectra before and after application of 10 µM Cu^2+^-TETAC in a fluorometer (see Methods and Approaches). As shown in Fig. 4, Cu^2+^-TETAC produced a large decrease in fluorescence intensity for both MBP-295Anap-C (Fig. 4A) and MBP-322Anap-C (Fig. 4B). The emission spectra in the presence of Cu^2+^ were nearly identical in shape to the spectra without Cu^2+^-TETAC (Fig. 4A-D, dashed traces) indicating the fluorescence quenching reflected a FRET mechanism as opposed to a change of environment of L-Anap or an inner filter effect.

**Fig. 4.**
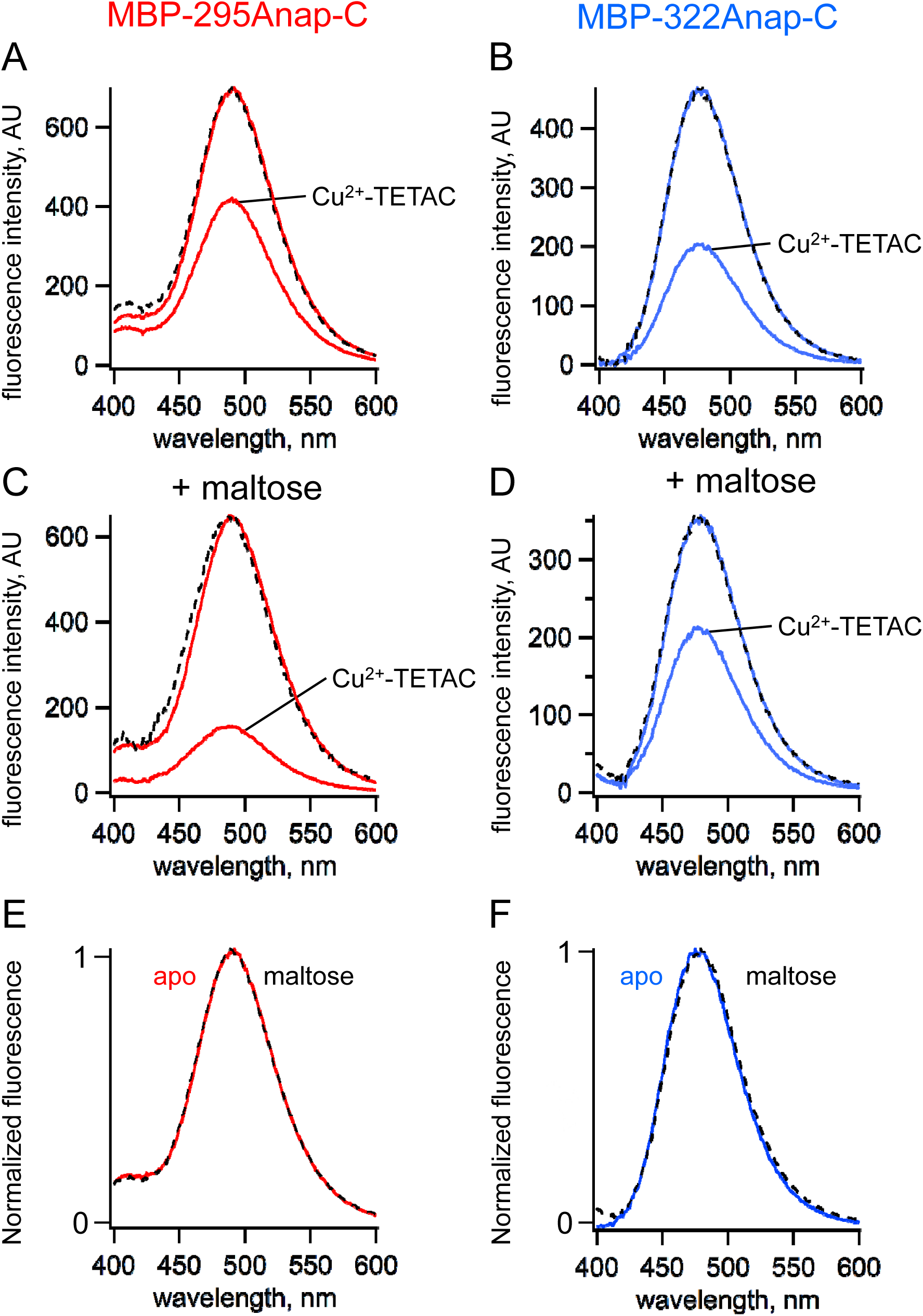
ACCuRET measures FRET and ligand-dependent changes in FRET. (A-B) Emission spectra of L-Anap for (A) MBP-295Anap-C and (B) MBP-322Anap-C measured in a fluorometer, before and after addition of 10 µM Cu^2+^-TETAC to the cuvette. The fluorescence quenching reveals FRET between L-Anap and Cu^2+^-TETAC. (C-D) Emission spectra of L-Anap for (C) MBP-295Anap-C and (D) MBP-322Anap-C, before and after addition of 10 µM Cu^2+^-TETAC, in the presence of 10 mM maltose. Maltose produced an increased FRET in MBP-295Anap-C and decreased FRET in MBP-322Anap-C. The dashed lines in panels A-D represent the Cu^2+^-TETAC spectra scaled to the initial spectra. The similarity indicates that quenching produced little or no change in the shape of the emission spectra, consistent with a FRET mechanism. (E-F) Scaled emission spectra of L-Anap for (E) MBP-295Anap-C and (F) MBP-322Anap-C with and without maltose. The similarity suggests that Anap experiences little or no environmental changes with maltose binding.

Fluorescence quenching was strongly dependent on the presence of maltose. For MBP-295Anap-C, the quenching was greater with maltose indicating an increase in FRET efficiency and shorter distance (Fig. 4C). For MBP-322Anap-C, the quenching was smaller with maltose indicating a decrease in FRET efficiency and longer distance (Fig. 4D). Changes in tmFRET in MBP-295Anap-C therefore allowed us to visualize the closing of the top of the clamshell with ligand binding, and changes in tmFRET in MBP-322Anap-C allowed us to visualize the opening of the backside of the clamshell (Supplemental Movie 1). Importantly, maltose did not change the shape of the emission spectra for either construct (Fig. 4E, F), indicating the environment of L-Anap did not change upon maltose binding.

The maltose affinity of wild-type MBP is about 2 µM (29). We found that wild-type MBP purified from HEK293T/17 cells appeared to be bound to an endogenous ligand, perhaps glycogen. It exhibited a FRET efficiency similar to the maltose-bound state, and did not exhibit the maltose-dependent change in tmFRET even after extensive washing and ion exchange chromatography (data not shown). Therefore, in all of the MBP constructs in this paper we introduced a mutation into the maltose binding site (W340A) that has previously been shown to reduce the maltose affinity (29). Indeed, this construct exhibited a maltose-dependent change in FRET (Fig. 4) indicating that did not copurify with an endogenous sugar.

We used ACCuRET to measure the maltose affinity of MBP-295Anap-C (containing W340A) modified with Cu^2+^-TETAC by measuring the fluorescence quenching as a function of maltose concentration. As shown in Fig. 5, maltose caused a concentration-dependent decrease in fluorescence with an apparent affinity of 280 µM, similar to the value measured previously with tryptophan fluorescence (29). No decease in fluorescence was observed in MBP-295Anap lacking the introduced cysteine (Fig. 5). These experiments illustrate the utility of ACCuRET for measuring both structural and functional properties of proteins.

**Fig. 5.**
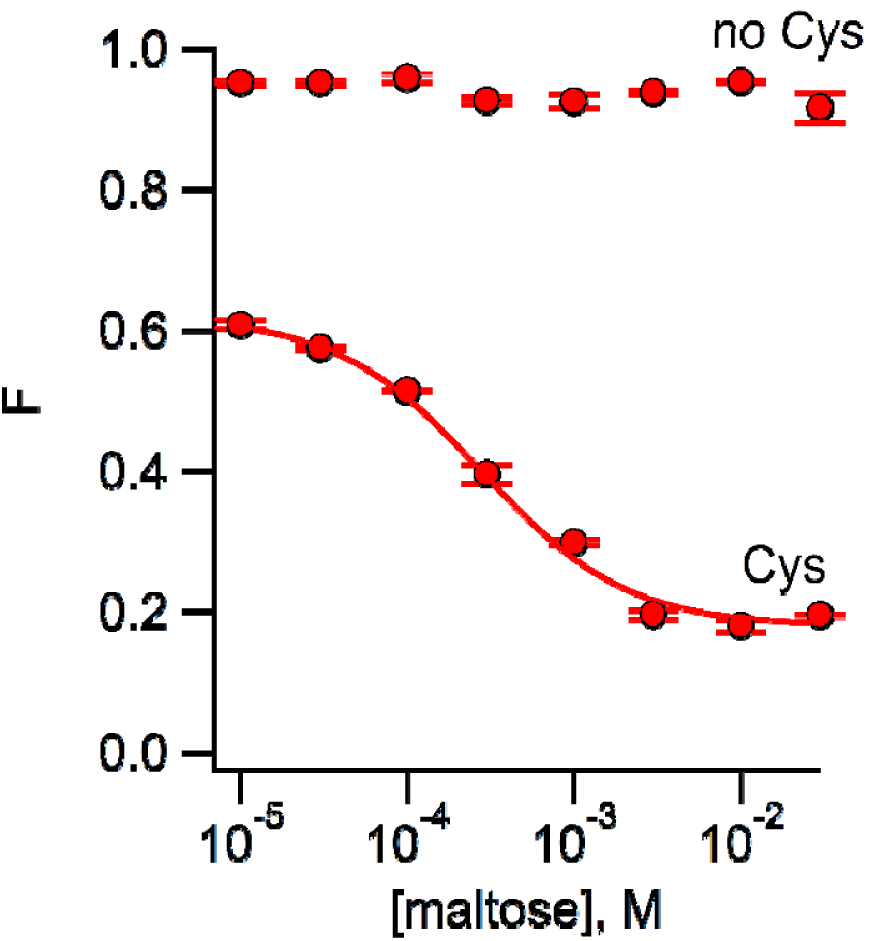
Using ACCuRET to measure the dependence of the rearrangement in MBP on maltose concentration. Plot of the fractional fluorescence of MBP-295Anap-C (Cys) and MBP-295Anap (no Cys), as indicated, after addition of Cu^2+^-TETAC, as a function of maltose concentration. Shown are mean ± SEM for n = 4. The smooth curve is a fit of a binding isotherm with an apparent affinity of 280 µM. The low affinity is expected from the W340A mutation we introduced into all of our MBP constructs to reduce binding to endogenous sugars (29).

To quantify ACCuRET, we measured the time course of the fluorescence at a given wavelength before and after addition of Cu^2+^-TETAC and the reducing agent DTT. For both MBP-295Anap-C and MBP-322Anap-C, there was a rapid (<10 s) and substantial quenching of the fluorescence upon addition of 10 µM Cu^2+^-TETAC (Fig. 6A, B). The quenching was nearly completely reversed upon addition of DTT (Fig. 6A, B) and negligible in the absence of the introduced cysteine (Fig. 6C, D), indicating that it resulted from a reaction of Cu^2+^-TETAC with the introduced cysteine. For MBP-295Anap-C the quenching was greater in the presence of maltose whereas for MBP-322Anap-C the quenching was less in the presence of maltose (Fig. 6C, D), as expected from the predicted distance changes. These results establish that ACCuRET can serve as a useful alternative to tmFRET with standard HH sites to create a minimal metal binding site for tmFRET with L-Anap and report intramolecular distance changes.

**Fig. 6.**
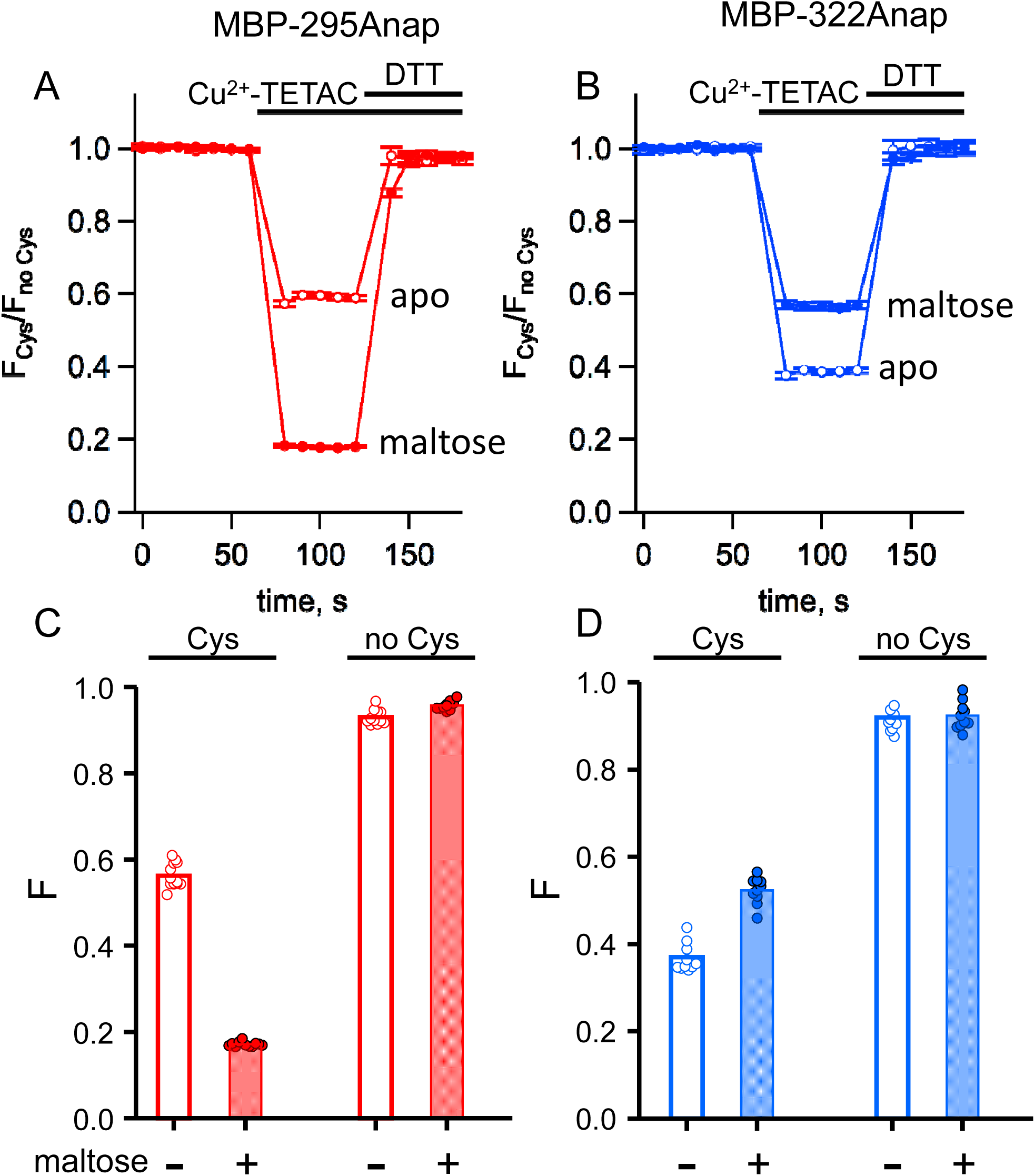
ACCuRET in MBP is reproducible, reversible, maltose dependent, and specific for each FRET pair. (A-B) Time course and reversal of ACCuRET for (A) MBP-295Anap-C and (B) MBP-322Anap-C in the absence (open symbols) and presence (filled symbols) of 10 mM maltose. The fractional fluorescence of each construct was recorded every 10 s and normalized to the fractional fluorescence of the corresponding construct without an introduced cysteine. 10 µM Cu^2+^-TETAC and 1 mM DTT were added to the cuvette at the times indicated by the bars. Shown are mean ± SEM for n = 6-8. (C-D) Scattergrams of fractional fluorescence of (C) MBP-295Anap and (D) MBP-322Anap, with (Cys) and without (no Cys) the introduced cysteine, after addition of 10 µM Cu^2+^-TETAC in the absence (-) and presence (+) of 10 mM maltose. The amount of quenching was reproducible, maltose dependent, construct dependent, and nearly absent without the introduced cysteine.

### tmFRET between L-Anap and transition metal bound to standard HH metal binding sites

We compared tmFRET efficiencies measured with ACCuRET to those measured with Cu^2+^ bound to standard HH sites, using the same donor-acceptor positions on MBP. Addition of 100 µM Cu^2+^ caused a rapid quenching of both MBP-295Anap-HH and MBP-322Anap-HH fluorescence (Fig. 7A, B). The quenching was nearly completely reversed upon addition of EDTA (Fig. 7A, B) and negligible in the absence of the introduced cysteine (Fig. 7C, D), indicating that it resulted from binding of Cu^2+^ to the introduced HH sites. For MBP-295Anap-HH, the quenching was greater in the presence of maltose whereas for MBP-322Anap-HH the quenching was less in the presence of maltose (Fig. 7), as observed with Cu^2+^-TETAC (Fig. 6).

**Fig. 7.**
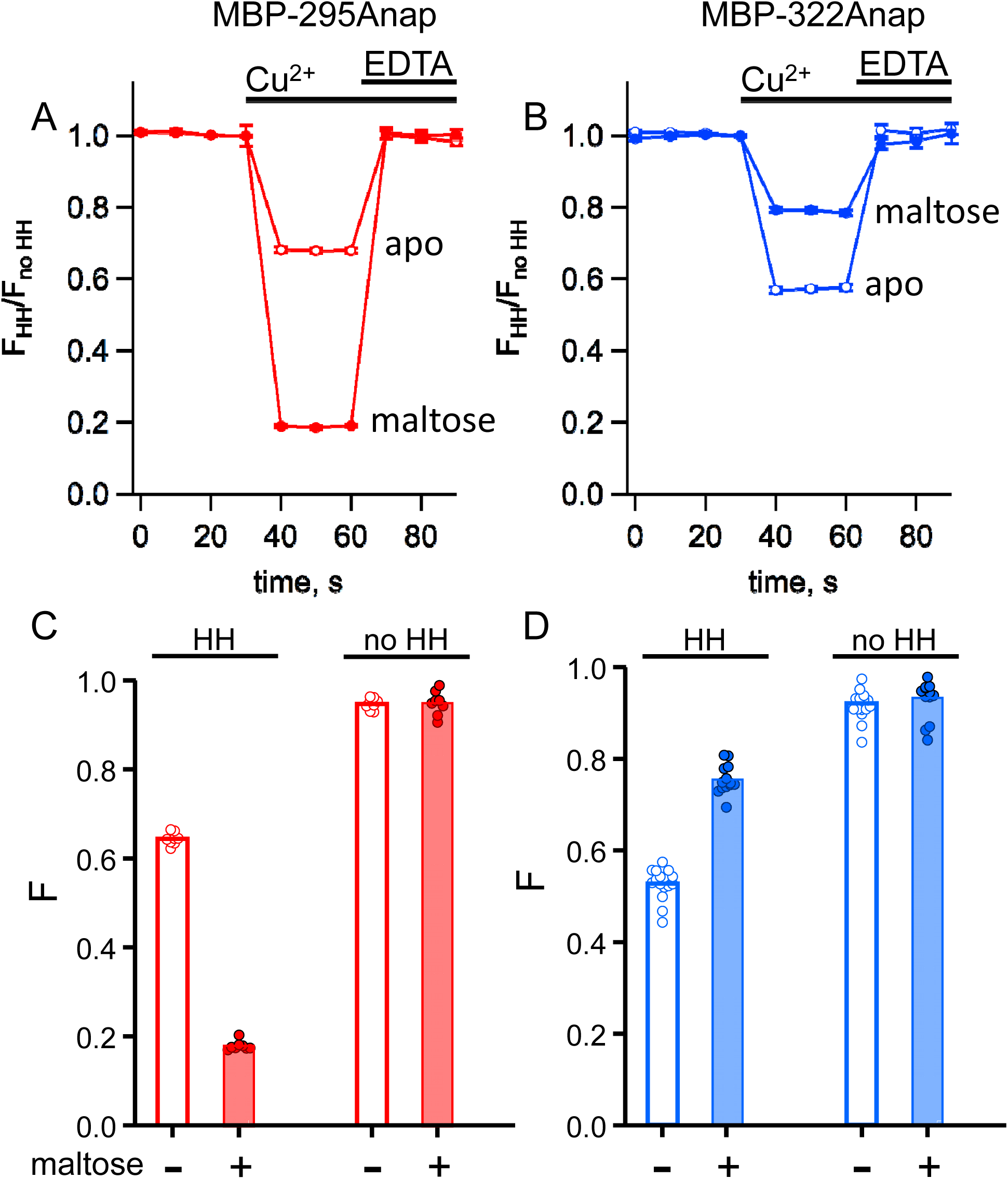
tmFRET with standard HH sites in MBP gives similar quenching to ACCuRET for the same FRET pairs. (A-B) Time course and reversal of tmFRET with HH sites for (A) MBP-295Anap-HH and (B) MBP-322Anap-HH in the absence (open symbols) and presence (filled symbols) of 10 mM maltose. The fractional fluorescence of each construct was recorded every 10 s and normalized to the fractional fluorescence of the corresponding construct without a HH site. 100 µM Cu^2+^ and 10 mM EDTA were added to the cuvette at the times indicated by the bars. Shown are mean ± SEM for n = 8. (C-D) Scattergrams of fractional fluorescence of (C) MBP-295Anap and (D) MBP-322Anap, with (HH) and without (no HH) the HH site, after addition of 100 µM Cu^2+^ in the absence (-) and presence (+) of 10 mM maltose. The amount of quenching with standard HH sites was similar to that with ACCuRET.

For both L-Anap sites, the quenching increased with increasing Cu^2+^ concentration, following a simple binding isotherm, with an apparent affinity corresponding to the affinity of the metal binding site (Supplemental Fig. 2). For Cu^2+^ binding to MBP-295Anap-HH (in the presence of maltose) the affinity was 3 µM (Supplemental Figure 2C), and for Cu^2+^ binding to MBP-322Anap-HH (in the absence of maltose) the affinity was 17 µM (Supplemental Figure 2D). The quenching at saturating concentrations reflects the FRET efficiency between L-Anap and the bound metal. These data support the use of 100 µM Cu^2+^ as a saturating concentration for measuring tmFRET in these constructs.

Interestingly, at 1 mM Cu^2+^ there was some fluorescence quenching in MBP-295Anap, but little in MBP-322Anap (Supplemental Fig. 2). As these constructs lack the HH metal binding site and 1 mM Cu^2+^ is a fairly high concentration, this quenching likely represents collisional quenching and suggests that 295Anap has a greater solvent accessibility compared to 322Anap, as also suggested by the emission spectra discussed above (Fig. 13B). These data also indicate that there was negligible quenching by Cu^2+^ bound to endogenous sites in MBP at Cu^2+^ concentrations below 1 mM. Although not an issue with the constructs used here, it is worth noting that these potential problems caused by high concentrations of free Cu^2+^ are largely mitigated in ACCuRET.

One advantage of standard HH sites is the ability to use transition metals with different coordination chemistries and absorption profiles than Cu^2+^ (Fig. 3C). These metals, including Co^2+^ and Ni^2+^, are expected to have different binding affinities for HH and endogenous sites as well as different R_0_ values. Co^2+^ produced a similar FRET efficiency to Cu^2+^ but exhibited about a 10-fold lower binding affinity (Supplemental Fig. 2C, pink). Ni^2+^, however, produced a substantially lower FRET efficiency at saturating concentration (Supplemental Fig. 2C, green). This reflects the lower R_0_ value predicted for Ni^2+^ and further supports a FRET mechanism for the energy transfer. More importantly, the lower R_0_ value for Ni^2+^ expands the useable range of tmFRET measurements to shorter distances. With standard HH sites, metals with different R_0_ values can be selected to match the distance of interest.

### Determination of distances and distance changes using ACCuRET

FRET efficiency is steeply dependent on distance between the donor and acceptor, making FRET a molecular ruler for measurements of distances (4). Accurate distance measurements, however, have historically been a challenge for FRET studies because they are highly dependent on the conformational heterogeneity and dynamics of the fluorophores, the relative orientation of the fluorophores, and the labeling specificity and efficiency (5, 6, 18). ACCuRET was designed to address these problems by using small probes with short linkers, a metal ion acceptor that is anisotropic, and labeling methods that are orthogonal, specific, and efficient.

Energy transfer from donor to acceptor causes quenching of donor fluorescence upon the addition of acceptor. With tmFRET, the FRET efficiency, E, can be easily calculated from the decrease in L-Anap fluorescence upon addition of metal acceptor. Other sources of energy transfer (e.g. solution quenching and off-target metal binding) and nonspecific decreases in fluorescence (e.g. bleaching and loss of protein) can confound calculations of FRET efficiency. The extent of this background quenching can be determined using constructs without an introduced metal binding site and the FRET efficiencies corrected for as described in Materials and Methods (8, 25, 26).

From the FRET efficiencies, we calculated the distances between the donors and acceptors using the Förster equation *R* = *R*_0_(1/*E* − 1)^1/6^ where *R*_0_ is the distance producing 50% FRET efficiency. We calculated *R*_0_ using the emission spectrum and quantum yield of L-Anap at each of the two positions in MBP, 295 and 322, and the absorption spectrum of Cu^2+^ bound to either a HH motif or cyclen. We assume random orientations of the donor and acceptor (*κ*^2^ = 2/3), a reasonable assumption when one member of the FRET pair is a metal ion (15).

ACCuRET accurately measured intramolecular distances and distance changes, comparable to tmFRET using standard HH sites. *Fig. 8* compares the tmFRET distance measurements for all our MBP constructs, with either Cu^2+^-TETAC or Cu^2+^-HH, in the absence (open circles) or presence (closed circles) of maltose. Also shown are the β-carbon distances predicted from the X-ray crystal structures of MBP in the absence and presence of ligand. Overall, the distances were highly accurate, with a root-mean-square deviation (RMSD) between measured distances and crystal structure distances of 1.5 Å for ACCuRET and 1.8 Å for tmFRET with standard HH sites. There was little variability in our measurements (Fig. 6C, D, and Fig. 7C, D), suggesting that the errors were more likely to be in our estimates of the probe distances from the β-carbon positions, or in assumptions of the Förster equation (see below), than in experimental variability due to the method. In addition, the changes in distances with maltose were similar in direction and magnitude to the changes in β-carbon distances predicted from the crystal structures. Finally, we did not detect a systematic difference between the distances measured with Cu^2+^-HH and Cu^2+^-TETAC (*Fig. 8*). In conclusion, ACCuRET accurately determined the absolute distances and maltose-dependent distance changes for the FRET pair at the top of the clamshell, for which the distance decreased with maltose, and the FRET pair on the back of the clamshell, for which the distance increased with maltose.

**Fig. 8.**
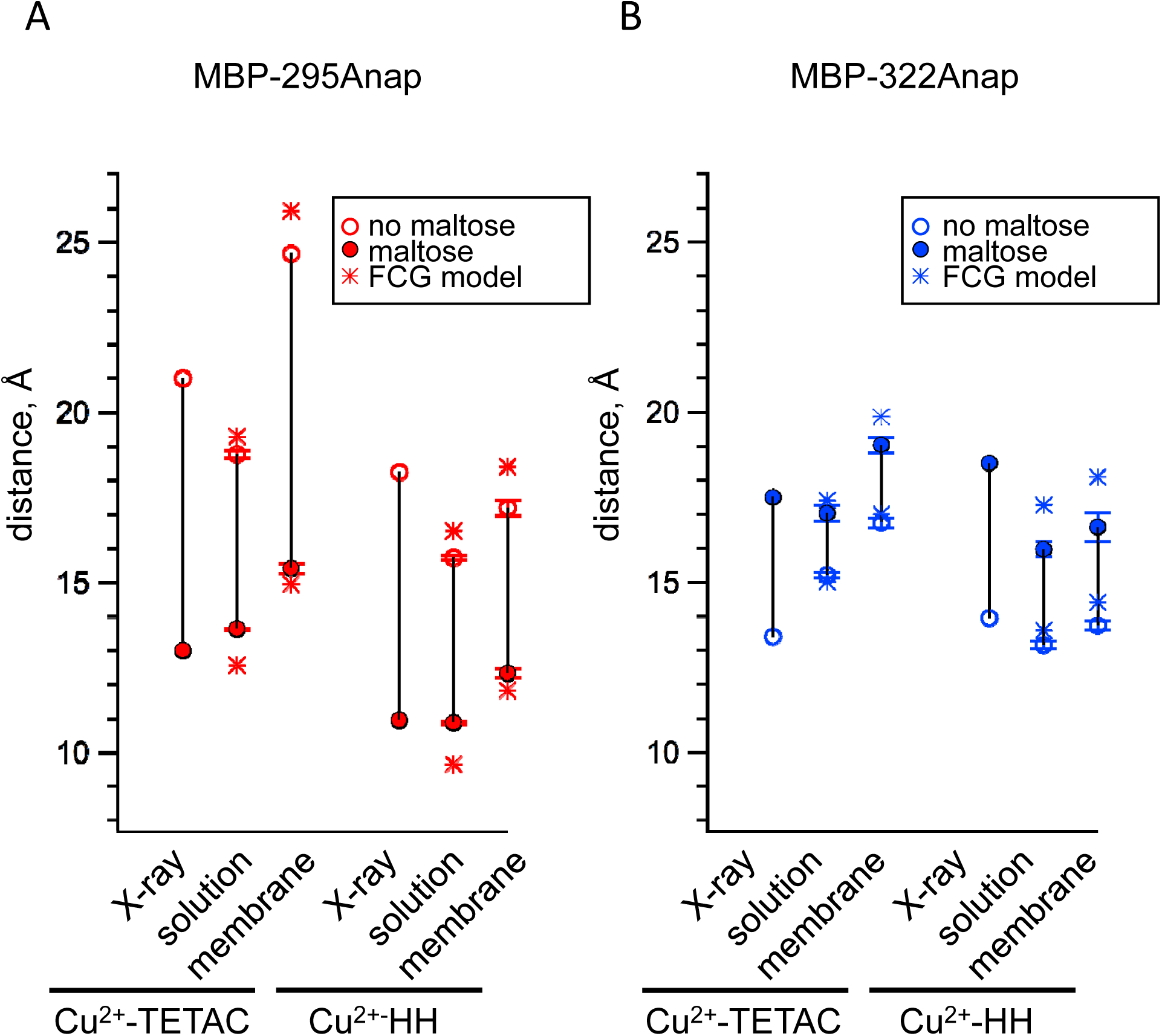
Comparison of distances measured with ACCuRET and standard HH sites to distances in the X-ray crystal structures of MBP (A-B) Distances from the X-ray crystal structures (X-ray) and determined using ACCuRET (Cu^2+^-TETAC) and standard HH sites (Cu^2+^-HH) from soluble (solution) and membrane-bound (membrane) MBP for the (A) MBP-295Anap and (B) MBP-322Anap FRET pairs. The distances determined using the Förster equation in the absence (open symbols) and presence (filled symbols) of maltose are shown and connected by a vertical line that reflects the maltose-dependent change in distance. Shown are mean ± SEM for n ≥ 8. The distances determined using the FCG model are shown as asterisks.

### ACCuRET measurements of distances in membrane proteins

Membrane proteins present a major challenge for measuring protein structure and conformational dynamics. Compared to soluble proteins, membrane proteins typically express at much lower levels, are difficult to purify in their native state, and often require their native membrane environment to exhibit physiological structural and functional properties. ACCuRET is well suited to address these challenges because it leverages the exquisite sensitivity of fluorescence measurements, the efficient incorporation of L-Anap into proteins in mammalian cells, the ability to work on unpurified proteins, and the ability to record from proteins in their native membranes. We therefore set out to validate and optimize ACCuRET for measuring conformational dynamics of proteins in the membrane.

To record fluorescence from native membranes we utilized a method called cell unroofing. Our implementation of this method utilizes a probe sonicator mounted on the stage of an inverted microscope to sheer off the dorsal surface of cells and dislodge all soluble cellular contents and organelles (*Fig. 9*; (17)). Cell unroofing leaves the ventral surface of cells intact as plasma membrane sheets attached to the coverslip for simple epifluorescence imaging. Cell unroofing also provides access to the intracellular surface of the membrane for application of transition metals and intracellular ligands. tmFRET measurements can then be made by recording the fluorescence from the unroofed cell before and after the application of Cu^2+^-TETAC or free transition metal.

**Fig. 9.**
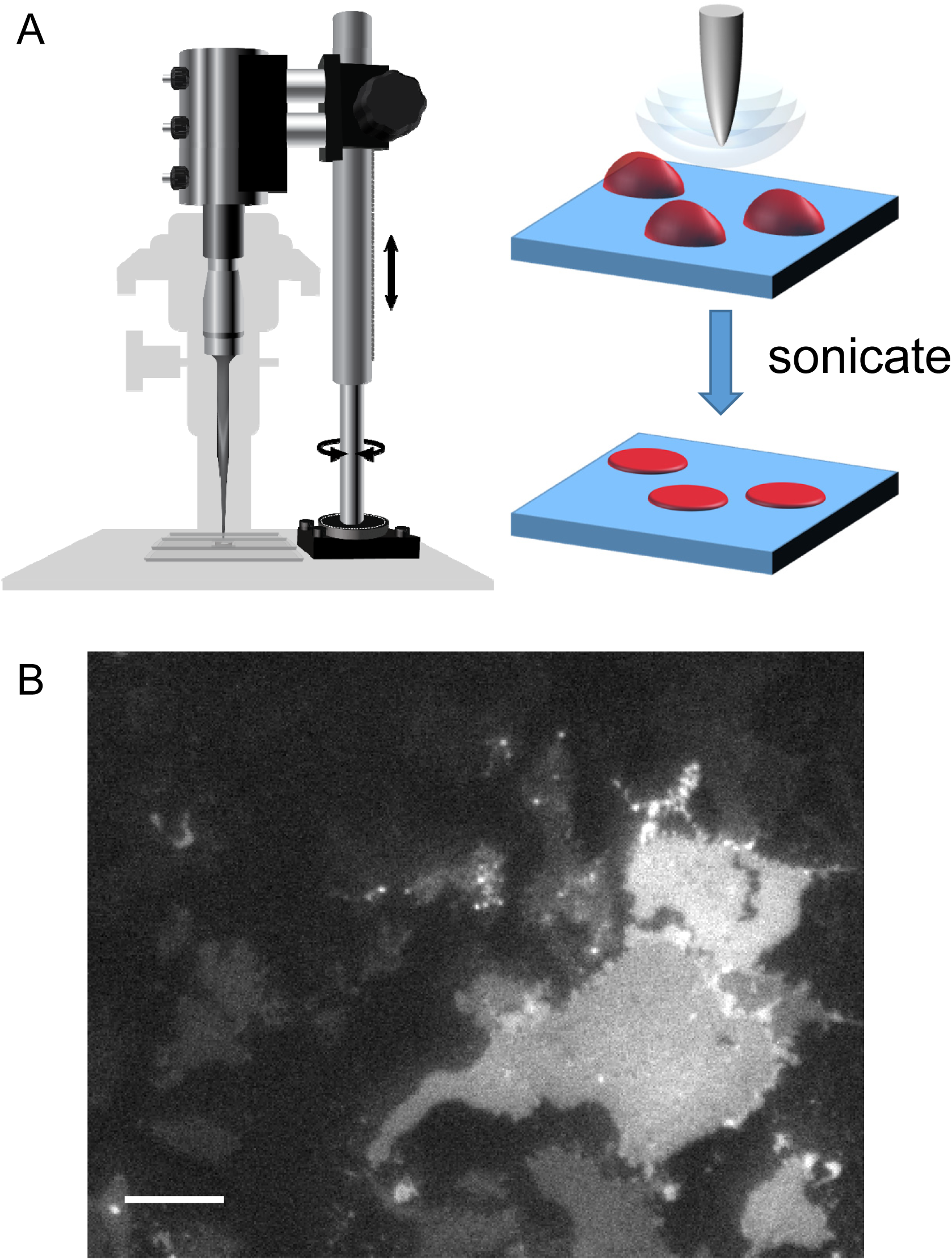
Cell unroofing to isolate plasma membrane sheets. (A) Diagram of the experimental setup used to unroof cells in a chamber on the microscope stage (left) with a cartoon of the unroofing process (right), as previously described (17). (B) An image of a field of unroofed HEK293T/17 cells transiently transfected with MBP-322TAG-CAAX and imaged with epifluorescence microscopy. The scale bar is 10 µm.

To validate and compare ACCuRET in unroofed cells, we made membrane-bound MBP constructs by adding a carboxy-terminal CAAX motif, which is then farnesylated in the cell (30, 31). These MBP-295TAG-CAAX and MBP-322TAG-CAAX constructs were each cotransfected into HEK293T/17 cells with pANAP and DN-eRF1 and incubated with L-Anap-ME in the medium, as described above. Both MBP-295Anap-CAAX and MBP-322Anap-CAAX expressed robustly and localized to the intracellular surface of the plasma membrane. Upon unroofing, the cells were no longer easily visible with bright field illumination. However, epifluorescence illumination showed plasma membrane sheets with relatively uniform L-Anap fluorescence (*Fig. 9*) which were stable for >30 minutes (data not shown). Previously, we showed that these images look similar with total internal reflection fluorescence (TIRF) microscopy, confirming that most or all of the fluorescence is from the plasma membrane (17). These results indicate that MBP-295Anap-CAAX and MBP-322Anap-CAAX proteins were stably associated with the membrane.

As for soluble MBP, Cu^2+^-TETAC was an effective tmFRET acceptor for membrane-bound MBP in unroofed cells. We unroofed the MBP-Anap-C-CAAX constructs and recorded the L-Anap fluorescence before and after addition of Cu^2+^-TETAC and DTT. As shown in *Fig. 10*A, C, Cu^2+^-TETAC produced a rapid decrease in L-Anap fluorescence of MBP-295Anap-C-CAAX that was reversed by the addition of DTT. The quenching was absent in MBP-295Anap-CAAX without the introduced Cys (*Fig. 10*E), indicating it arose from Cu^2+^-TETAC modification of this Cys. The quenching was greater in the presence of maltose (*Fig. 10*C, E) as expected from the decrease in distance between the donor and acceptor with maltose for the 295 FRET pair (Supplemental Movie 1). Similarly, for MBP-322Anap-C-CAAX, Cu^2+^-TETAC produced a rapid and reversible decrease in L-Anap fluorescence that was absent in MBP-322Anap-CAAX (*Fig. 10*B, D). With MBP-322Anap-C-CAAX, however, the quenching was less in the presence of maltose (*Fig. 10*D, F), as expected from the maltose-induced increase in distance between the donor and acceptor for the 322 FRET pair (Supplemental Movie 1). The fractional fluorescence quenching was highly reproducible from cell to cell and was similar to the results obtained for soluble MBP in a cuvette (Fig. 6). These results establish that ACCuRET can be used to measure tmFRET in membrane proteins.

**Fig. 10.**
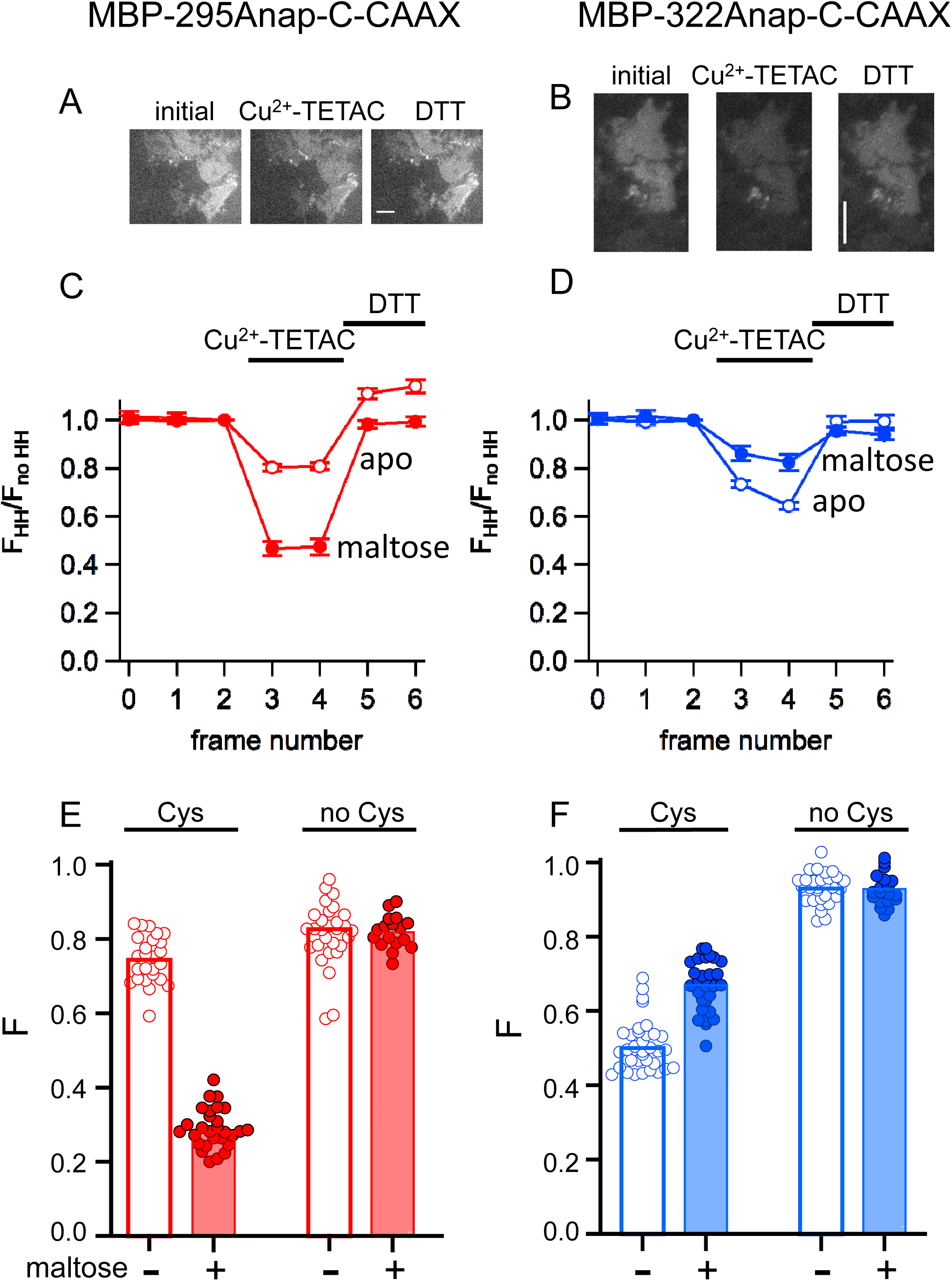
ACCuRET in membrane-bound MBP in unroofed cells (A-B) Images of unroofed cells expressing (A) MBP-295Anap-C-CAAX or (B) MBP-322Anap-C-CAAX showing quenching upon addition of 10 µM Cu^2+^-TETAC and reversal upon addition of 1 mM DTT. The scale bar is 10 µm. Time course and reversal of ACCuRET for (A) FLAG-MBP-295Anap-C-CAAX and (D) FLAG-MBP-322Anap-C-CAAX in the absence (open symbols) and presence (filled symbols) of 10 mM maltose. The fractional fluorescence of each construct was recorded every 10 s and normalized to the fractional fluorescence of the corresponding construct without an introduced cysteine. 10 µM Cu^2+^-TETAC and 1 mM DTT were added to the chamber at the times indicated by the bars. Shown are mean ± SEM for n ≥ 17. (E-F) Scattergrams of fractional fluorescence of (E) FLAG-MBP-295Anap-C-CAAX, and (F) FLAG-MBP-322Anap-C-CAAX, with (Cys) and without (no Cys) the introduced cysteine, after addition of 10 µM Cu^2+^-TETAC in the absence (-) and presence (+) of 10 mM maltose. The amount of quenching was reproducible, maltose dependent, construct dependent, and nearly absent without the introduced cysteine.

We compared tmFRET efficiencies measuring with ACCuRET to those measured with Cu^2+^ bound to standard HH sites, using the same donor-acceptor positions. Addition of 100 µM Cu^2+^ to unroofed cells expressing MBP-295Anap-HH-CAAX and MBP-322Anap-HH-CAAX produced a rapid decrease in fluorescence that was reversed by EDTA and nearly absent without the introduced HH sites (*Fig. 11*). Furthermore, the quenching with Cu^2+^ was greater with maltose in MBP-295Anap-HH-CAAX and less with maltose in MBP-322Anap-HH-CAAX. The fractional fluorescence quenching was highly reproducible from cell to cell and closely mirrored the results obtained for soluble MBP in a cuvette.

**Fig. 11.**
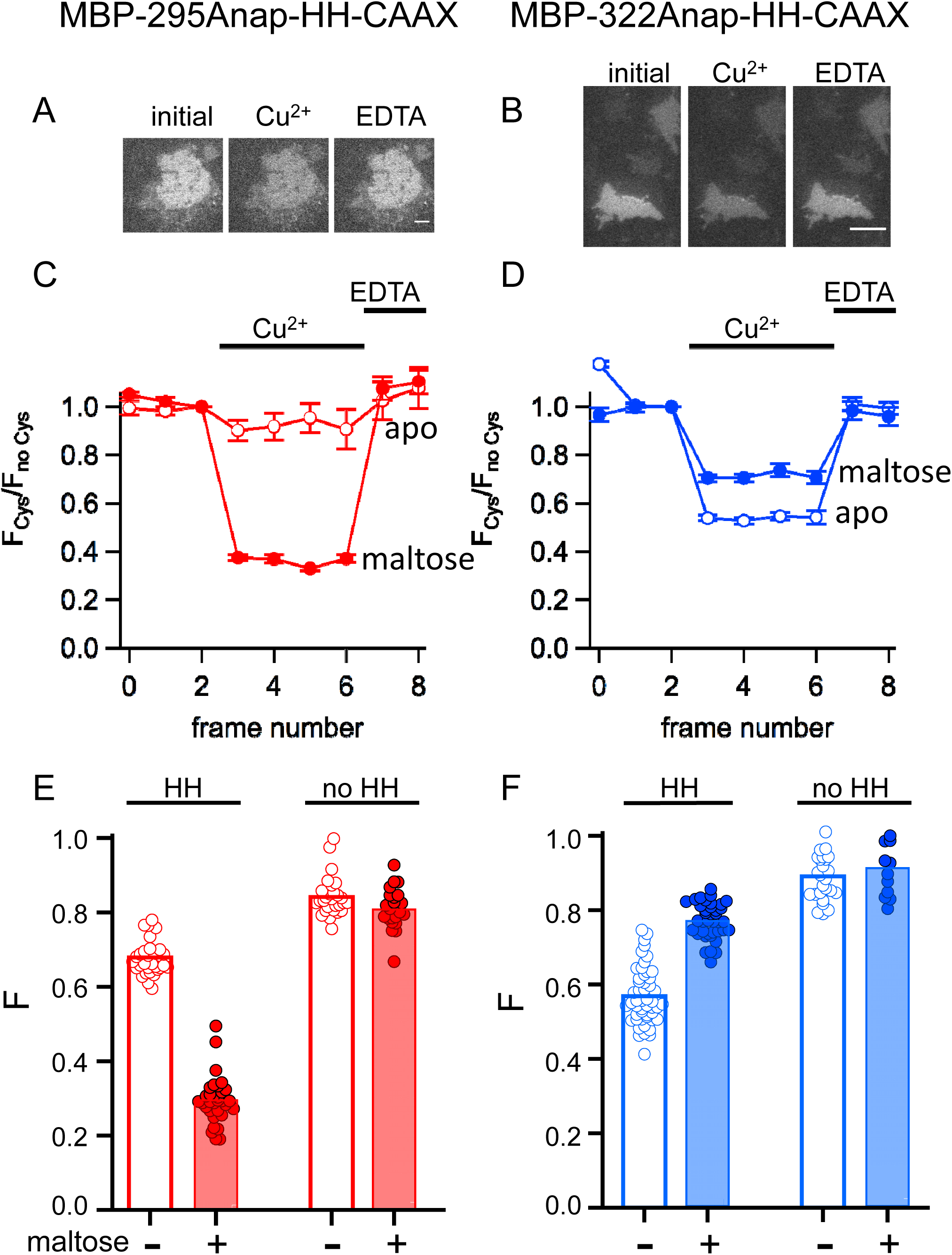
tmFRET with HH sites in membrane-bound MBP in unroofed cells (A-B) Images of unroofed cells expressing (A) MBP-295Anap-HH-CAAX or (B) MBP-322Anap-HH-CAAX showing quenching upon addition of 100 µM Cu^2+^ and reversal upon addition of 10 mM EDTA. The scale bar is 10 µm. (C-D) Time course and reversal of tmFRET with HH sites for (C) MBP-295Anap-HH-CAAX or (D) MBP-322Anap-HH-CAAX in the absence (open symbols) and presence (filled symbols) of 10 mM maltose. The fractional fluorescence of each construct was recorded every 10 s and normalized to the fractional fluorescence of the corresponding construct without a HH site. 100 µM Cu^2+^ and 10 mM EDTA were added to the chamber at the times indicated by the bars. Shown are mean ± SEM for n ≥ 12. (E-F) Scattergrams of fractional fluorescence of (E) MBP-295Anap-HH-CAAX or (F) MBP-322Anap-HH-CAAX, with (HH) and without (no HH) the HH site, after addition of 100 µM Cu^2+^ in the absence (-) and presence (+) of 10 mM maltose. The amount of quenching with standard HH sites was similar to that with ACCuRET.

From the FRET efficiencies, we calculated the distances between the donors and acceptors using the Förster equation as described above. Comparing our determination of distances from membrane-bound MBP to those predicted from the X-ray crystal structures of MBP (*Fig. 8*) gives RMSD values of 2.9 Å for ACCuRET and 1.3 Å for tmFRET with standard HH sites, comparable to the values for MBP in solution. However, distances calculated from membrane-bound MBP-Anap constructs were systematically longer than those determined from MBP-Anap constructs in solution (see Discussion). Together, our data demonstrate that ACCuRET provides accurate determinations of distances and distance changes in both soluble proteins and membrane proteins.

### Heterogeneity

The Förster equation predicts a steep distance dependence for FRET efficiency (*Fig. 12*A, black curve) and assumes that the donor and acceptor probes are a fixed distance apart (*Fig. 12*B, black). However, the linkers between the probes and the protein backbone produce substantial heterogeneity and mobility of the probes that effect the measured FRET efficiency (5, 6, 9). This heterogeneity in distance, together with the nonlinear dependence of the FRET efficiency on distance, cause the distances closer to the *R*_0_ distance to be weighted more heavily. The distance dependence of FRET efficiency, therefore, becomes shallower than predicted by the Förster equation. Our approach to reduce this effect has been to minimize these linkers by using a fluorescent amino acid and minimal metal binding sites closely associate with the backbone (HH or TETAC). Still, we observed a small, systematic bias in our tmFRET measurements where the distances greater than *R*_0_ have somewhat greater FRET efficiency and the distances less than *R*_0_ have somewhat lower FRET efficiency than predicted by the Förster equation (*Fig. 12*A). This manifests as an apparent shallowing of the distance dependence and an underestimation of the changes in distance with maltose.

**Fig. 12.**
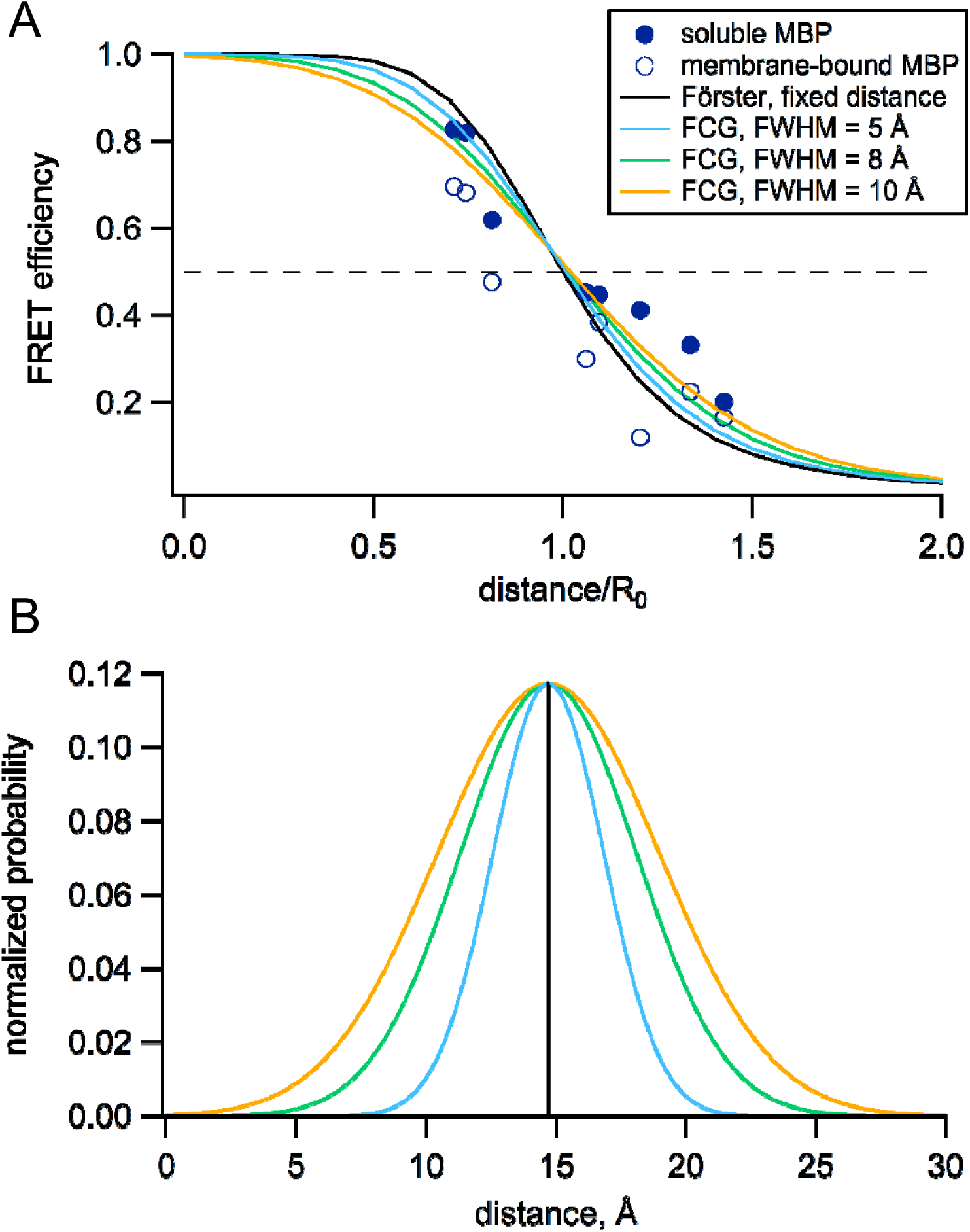
Fӧrster equation convolved with Gaussian distribution to predict average distances. (A) Plot of the measured and predicted distance dependence of FRET efficiency. The FRET efficiencies measure in this paper from soluble (filled symbols) and membrane-bound (open symbols) MBP are plotted versus the Cβ-Cβ distances predicted from the MBP X-ray crystal structures (PDB IDs: 1N3W and 1N3X for apo and maltose-bound, respectively). The predicted distance dependencies are show for the Fӧrster equation and the Fӧrster equation convolved with Gaussian distribution. The Gaussians used for convolution had mean intramolecular distances of 14.7 Å and the following FWHM values: 0 Å (black), 5 Å (blue), 8 Å (green), and 10 Å (orange). (B) The Gaussian distributions that were convolved with the Fӧrster equation for panel (A), using the same color scheme.

To correct for this heterogeneity, we developed a method we call FCG (<u>F</u>örster <u>c</u>onvolved with <u>G</u>aussian) analysis. FCG analysis assumes a Gaussian distribution of donor-acceptor distances in the sample (Fig. 12B). This distribution is typical of the distance distributions between two spin labels on rigid proteins measured using double electron-electron resonance (DEER) spectroscopy (32). When convolved with the Förster equation, the apparent FRET efficiency has a similar (but not identical) *R*_0_ but a more shallow distance dependence than the Förster equation (*Fig. 12*A). The greater the width of the Gaussian distribution, the shallower the distance dependence. Using a Gaussian distribution with a full width at half maximum (FWHM) of 8 Å, we found that, in most cases, the distances and changes in distance calculated from the FCG model (*Fig. 8*, asterisks) more closely matched the distances predicted from the crystal structures than the distances calculated with the Förster equation alone. Since all proteins at physiological temperature possess some structural heterogeneity (33), we propose that this correction may be useful for many FRET studies of distance.

## Discussion

In this paper, we introduced a new method, called ACCuRET, for measuring intramolecular distances and dynamics in proteins using tmFRET between a donor fluorophore and an acceptor transition metal ion. For the donor fluorophore, ACCuRET utilizes a fluorescent noncanonical amino acid, L-Anap, site-specifically incorporated into the protein using amber codon suppression. Orthogonal labeling with metal acceptor was achieved by modifying an introduced cysteine with Cu^2+^-TETAC. Using MBP as a benchmark, we show that ACCuRET accurately measures absolute distances and changes in distance that faithfully represent the backbone dynamics underlying function in soluble and membrane proteins. Measurements of distance display an RMSD of 1.5-2.9 Å between our experimentally-determine distances and the X-ray crystal structures. These experiments validate ACCuRET as a powerful new method for measuring structural dynamics of proteins in their native environment.

The noncanonical amino acid L-Anap has been previously used to label membrane proteins. However, most of these studies focused on state-dependent changes in fluorescence due to the environmentally-sensitive emission of L-Anap (21, 24, 25, 34-37). L-Anap has also been recently used as a FRET donor in studies to discern movement in ion channels between domains or between the channel and the membrane (17, 22, 24, 25). Although these studies demonstrated the potential power of using L-Anap as a donor for tmFRET, the distances and distance changes measured were unexpected and difficult to interpret, given the limited structures available. By using MBP as a benchmark, we have been able to assess the accuracy of tmFRET for measurements of distances and changes in distance.

The use of amber codon suppression to introduce noncanonical amino acids has become more common in prokaryotic expression systems, but has been limited in eukaryotic expression systems by the much poorer efficiency of incorporation observed. We found that coexpression of a plasmid encoding our TAG-containing MBP with one encoding the DN-eRF1 was critical in achieving sufficiently high levels of expression (23). In addition to increasing the total amount of protein, DN-eRF1 also enhanced the fraction of the protein that was full-length. Thus, DN-eRF1 extends the power of amber codon suppression to mammalian cells and is particularly useful for eukaryotic membrane proteins that typically cannot be expressed in bacteria.

ACCuRET uses a novel metal-labeling strategy for tmFRET that is complementary, and in some ways superior to, the standard method of labeling via HH sites. Cu^2+^-TETAC has previously been used in paramagnetic relaxation enhancement (PRE) and DEER experiments (38, 39). The DEER experiments showed that it has a narrow inter-label distance distribution, very similar to that of the widely-used spin label MSL and much narrower than MTS-EDTA (38). TETAC labels cysteines in any secondary structure and has a subnanomolar affinity for Cu^2+^, allowing ACCuRET to be implemented in a wider variety of conditions. Native cysteines are mostly compatible with the use of TETAC. Only native cysteines within ~20 Å of the L-Anap, or for which modification produces functional effects, are problematic. Such native cysteines can easily be identified using the control construct without the introduced cysteine, and only those that are problematic need be mutated. Because the affinity of Cu^2+^ for TETAC is so high, Cu^2+^-TETAC can be rinsed from the bath solution prior to imaging unroofed cells, allowing measurements to be made in the absence of free Cu^2+^. Labeling with Cu^2+^-TETAC can be reversed with a reducing agent such as DTT, obviating the need for a metal chelator such as EDTA that can strip off endogenous metals and divalent ions. Although we expected the position of Cu^2+^ to be less constrained when bound to TETAC compared to HH, our distance measurements were as accurate for Cu^2+^-TETAC as for Cu^2+^-HH (*Fig. 8*). Finally, binding to the cyclen ring of TETAC increases the absorption of Cu^2+^ and blue shifts its spectrum (Fig. 3C), Cu^2+^-TETAC has an R_0_ value almost 4 Å longer than Cu^2+^ bound to HH (Fig. 3D). This longer R_0_ value increases the range of distance measurements that can be accurately measured with tmFRET.

Previously, the same donor/acceptor sites on MBP we use here have been studied with tmFRET using standard methods of labeling (7). At the donor site, the fluorophore fluorescein-5-maleimide or monobromobimane was reacted with an introduced cysteine, and, at the acceptor site, Cu^2+^ or Ni^2+^ was bound to a HH site. For the same donor/acceptor sites, our measurements of distances and changes in distance with ACCuRET were comparable with the measurements made using bimane as the donor, but more accurate than the measurements made using fluorescein as the donor, emphasizing the need for small donor fluorophores with short linkers. Importantly, however, these previous experiments required the use of purified protein with no native cysteines or metal binding sites. By employing a fluorescent noncanonical amino acid as the donor, ACCuRET can be performed on unpurified protein in its native environment with most or all of the native cysteines and weak metal binding sites left intact.

The dearth of approaches for studying conformational dynamics of proteins in their native environment makes ACCuRET especially valuable for studies of membrane proteins such as ion channels and transporters. We therefore compared ACCuRET in membrane-bound MBP in unroofed cells to soluble, affinity-purified MBP. Overall, the distance measurements in the membrane-bound MBP were similar to those in soluble MBP (*Fig. 8*). We observed, however, a trend toward slightly lower FRET efficiency for membrane-bound MBP than observed with soluble MBP (*Fig. 12*). The most likely explanation for this difference, which translated into ~2-3 Å increase in distance for our measurements in membrane-bound MBP compared to soluble MBP, is that our background subtraction method (see Materials and Methods) did not fully account for all the L-Anap-dependent background signal. Determining the source of the background fluorescence and developing better corrections will further improve the accuracy of ACCuRET for membrane proteins and, perhaps, proteins in intact cells.

ACCuRET mitigates a number of limitations that have hampered measurements of FRET efficiencies and calculations of absolute distances. Because ACCuRET measures L-Anap fluorescence in the absence and presence of the acceptor for both the apo and ligand-bound states, it accounts for any ligand-dependent changes in environment that would otherwise affect the calculation of FRET efficiency. Our measurements of the emission spectra of L-Anap incorporated at the two sites in MBP allowed us to use the R_0_ determined for each site in our calculations of distance. The difference in R_0_ between MBP-295Anap (17.7 Å) and MBP-322Anap (16.5 Å), however, was small and did not substantially affect our estimates of distances or changes in distance. The difference in quantum yield between our two sites (MBP-295Anap:0.31 and MBP-322Anap:0.47) is larger than generally observed for the state-dependent changes in fluorescence at any given site, suggesting that state-dependent changes in R_0_ are not always a concern.

Experimental limitations that could compromise the accuracy of distance determinations include: a broad distance distribution between donor and acceptor; measuring distances outside the range of 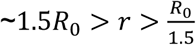 (Fig. 12B); nonspecific labeling with the donor; incomplete labeling with the acceptor; and unexpected sources of background fluorescence. ACCuRET mostly overcomes these limitations. Using small probes with short linkers narrows the distance distribution, and FCG analysis improves distance determinations for distributed distances. The increase in absorption of Cu^2+^-TETAC, and the corresponding increase in R_0_ values, expands the utility of tmFRET over a broader distance range. Using amber codon suppression to introduce the donor largely eliminates nonspecific labeling with donor. Using a slight excess of Cu^2+^ with TETAC minimizes incomplete labeling with the acceptor. As discussed above, unexpected sources of background fluorescence appear to reduce the accuracy of distance measured for membrane-bound MBP, but this effect was small in our experiments.

The distance dependence of FRET is, in practice, less steep than predicted by the Fӧrster equation (*Fig. 12*A). This is at least partly explained by the idealized assumption of the Fӧrster equation that relative distances between donors and acceptors are homogeneous (5, 6). In fact, proteins have been shown to exhibit significant heterogeneity (33), with distances between side-chains well described by normal distributions (32). Convolving Gaussian distance distributions with the Fӧrster equation (FCS analysis) gives distance-dependence curves that are less steep than the Fӧrster equation itself (*Fig. 12*A). From DEER studies, the distribution of distances between Cu^2+^ ions bound to TETAC were well described by Gaussian distributions with FWHM values ranging from 6 to 9 Å (38). We used FCG analysis to convert FRET efficiencies to distances. These distances (*Fig. 8*, asterisks) more closely match the donor-acceptor distances, as well as the maltose-induced distance changes, determined from the Cβ-Cβ values from X-ray crystal structures. Although we did not measure the distributions of donor-acceptor distance in our experiments, it seems clear that assuming a distribution of distances in the range found in the literature is a better assumption than assuming a fixed distance.

In summary, these experiments establish a new method called ACCuRET for measuring structural dynamics of proteins in their native environment, particularly membrane proteins. The method can measure distances with an accuracy of 1.5-2.9 Å and has the potential to measure structural dynamics on a time scale of milliseconds. For ion channel and transporters, ACCuRET can also be combined with patch-clamp fluorometry (PCF) to measure protein structure and function simultaneously. Although we used the unnatural amino acid L-Anap, our approach could employ fluorophores introduced with other unnatural amino acids (perhaps called unACCuRET). Ultimately, better fluorophores will enable tmFRET measurements with faster time resolution and single-molecule sensitivity.

## Materials and Methods

### Constructs, cell culture, and transfection

All constructs were made in the pcDNA3.1 mammalian expression vector (Invitrogen, Carlsbad, CA), except as noted below. The amino acid sequence of MBP was based on the pMAL-c5x vector (New England Biolabs, Ipswich, MA), and was codon optimized for mammalian cell expression. To reduce the affinity of MBP for endogenous ligands and allow subsequent testing of maltose binding, we introduced a point mutation, W340A (29), in all constructs. In addition, a FLAG epitope was added to the N-terminus of MBP. FLAG-MBP cDNA was synthesized by Bio Basic (Amherst, NY). The FLAG epitope is present in all constructs, but is omitted from the construct names except in the discussion of Western blot analysis. For membrane localization, the C-terminus of FLAG-MBP was fused to a CAAX domain with the following sequence: KMSKDGKKKKKKSKTKCVIM. The pAnap vector ((16); purchased from Addgene, Cambridge, MA) was used as previously described (17). The dominant negative eukaryotic release factor 1 construct (DN-eRF1) was kindly provided by Jason Chin (Cambridge, UK) and used in the provided pcDNATM5/FRT/TO vector (23).

Amber stop codons (TAG), histidines and cysteines were introduced using standard PCR and oligonucleotide-based mutagenesis. All sequences were confirmed using automated DNA sequencing (Eurofin/Operon, Louisville, KY). The stop codons were introduced into FLAG-MGP at positions 295 (MBP-295TAG) and 322 (MBP-322TAG). For each stop codon, three constructs were produced: no metal binding site, a di-histidine, and a cysteine. For MBP-295TAG, the di-histidines were introduced at positions 233 and 237 (MBP-295TAG-HH), and the cysteine was introduced at position 237 (MBP-295TAG-C). MBP-295TAG-C also contained the 233H mutation. For MBP-322TAG, the di-histidines were introduced at positions 305 and 309 and the cysteine at position 309.

All experiments utilized HEK293T/17 cells plated in 6-well trays. Cells were transfected at ≈25% confluency with a total of 1.6 µg of DNA and 10 µL of Lipofectamine 2000 (Invitrogen, Carlsbad, CA) per well. The 1.6 µg of DNA consisted of 0.9 µg of FLAG-MBP, 0.3 µg of pANAP and 0.4 µg of DN-eRF1. For experiments in which DN-eRF1 was not used, it was replaced with 0.4 µg pcDNA3. The DNA/Lipofectamine mix was prepared in 300 µL Opti-MEM (Invitrogen, Carlsbad, CA) per well. For transfection, cells were incubated in growth medium without antibiotics for 4 – 6 hours at 37 °C with 5% CO_2_. After incubation, the medium was replaced with one including antibiotics and supplemented with 20 µM L-Anap-ME (AssisChem, Waltham, MA). The L-Anap-ME was made as a 10 mM stock in ethanol and stored at −20 °C. Trays were wrapped with aluminum foil to block any light and incubated at 37 °C until use.

For fluorometry or Western blot experiments, cells were harvested approximately 24 hours after transfection. Cells were washed twice with PBS, and cell pellets were stored at −20 °C until use. For imaging experiments, the day after transfection medium was removed from the wells and replaced with HEPES buffered Ringers (HBR) solution (in mM: NaCl 140; KCl 4; CaCl_2_ 1.8; glucose 5; HEPES 10; pH 7.4). Cells were then placed on the bench top at room temperature for the remainder of the same day.

### Protein purification and Western blot analysis

Between 1 and 4 frozen pellets were thawed and resuspended in 0.5-0.8 mL Stabilization Buffer Tris (SBT) (in mM: KCl 70; MgCl_2_ 1; Trizma Base 30; pH 7.4) supplemented with cOmplete mini EDTA-free protease inhibitor cocktail (Sigma-Aldrich, St. Louis, MO). The suspension was sonicated using a Sonifier 450 with MicroTip (Branson, Danbury, CT) with settings of power = 4 and duty cycle = 50% for a total of 10 pulses. Lysed cells were then spun in a benchtop centrifuge at 13,000 rpm for 20 minutes, and the cleared lysate was moved to a new tube.

Anti-FLAG M2 affinity gel (Sigma-Aldrich, St. Louis, MO) was prepared by rinsing 100 µL of slurry per sample five times with 1 mL SBT. Cleared lysate was then added to the rinsed gel and nutated at 4 °C for one to two hours. Tubes were wrapped in aluminum foil to prevent photodamage. The gel was rinsed five times with 1 mL of SBT. Between 0.25 to 0.5 mL of a 200 ng/mL solution of FLAG peptide (Sigma-Aldrich, St. Louis, MO) in SBT was added to the rinsed beads, which were then nutated at 4 °C for between 90 minutes and 12 hours to elute the protein. The purified protein was harvested by spinning the beads and collecting the supernatant. The protein was stored in tubes wrapped in aluminum foil at 4 °C for up to one week.

For Western blot analysis, proteins were run on NuPage 10% bis-tris gels (ThermoFisher Scientific, Waltham, MA) with MOPS SDS running buffer (1 M MOPS, 1 M Tris, 69.3 mM SDS and 20.5 mM EDTA). Proteins were transferred to PVDF membranes using a BioRad Transblot SD (Hercules, CA) transfer cell with Bjerrum/Schafer-Nielsen transfer buffer with SDS (40). Membranes were blocked in 5% Milkman Instant Low Fat Dry Milk (Marron Foods, Harrison, NY) in TBS-T (20 mM Tris, 137 mM NaCl, 0.1% Tween20, pH 7.6) for either 1 hour at room temperature or overnight at 4 °C. Anti-FLAG primary antibody (Sigma-Aldrich Cat F3165, St. Louis, MO) was used at a dilution of 1:20,000 and secondary antibody (Amersham Cat NA931, Pittsburgh, PA) HRP-linked anti mouse IgG was used at 1:30,000 dilution in TBS-T. Secondary antibodies were visualized with Super Signal West Femto Substrate (ThermoFisher, Waltham, MA) and imaged with a Proteinsimple gel imager (San Jose, CA).

### Fluorometry and spectrophotometry

Starna (Atascadero, CA) sub-micro fluorometer cells (100 µL) were used for both fluorometry and spectrophotometry. Fluorometry experiments were performed using a Jobin Yvon Horiba FluoroMax-3 spectrofluorometer (Edison, NJ). For emission spectra of L-Anap, we used an excitation wavelength of 370 nm and 5 nm slits for excitation and emission, except for experiments to measure quantum yield in which 1 nm slits were used. For time course measurements, we excited samples at 370 nm and recorded the emission at 480 nm every ten seconds using anti-photobleaching mode of the instrument. Reagents (transition metals, Cu^2+^-TETAC, and DTT) were added manually as 100x stocks during the interval between measurements by pipetting up and down in the cuvette without removing it from the instrument.

Protein samples were diluted 1:10 to 1:200 in SBT to keep the fluorescence intensity within the linear range of the spectrophotometer. We found that it was essential to use Tris buffered solutions for experiments with Cu^2+^ and Cu^2+^-TETAC. Transition metal ions (Cu^2+^, Co^2+^ and Ni^2+^) were prepared from sulfate or chloride salts as 100 mM stocks in water or SBT, with stocks of lower concentrations diluted from this stock in SBT. TETAC (Toronto Research Chemicals, Toronto, Canada) was prepared as a 100 mM stock in DMSO and stored at −20 °C until the day of use. The TETAC stock appeared stable over a time scale of months and tolerated multiple freeze-thaw cycles. To prepare Cu^2+^-TETAC, 1 µL each of 100 mM TETAC stock and 110 mM CuSO_4_ stock were mixed together and allowed to incubate for one minute as the solution turned a deeper shade of blue, reflecting binding of Cu^2+^ to the cyclen ring. To this mixture, 98 µL of SBT was added, giving a solution of 1.1 mM Cu^2+^ and 1 mM TETAC. The high concentrations in the binding reaction and 10% over-abundance of Cu^2+^ ensured that all of the TETAC was bound with Cu^2+^. This stock solution was then diluted 1:100 when added to the cuvette for fluorometer experiments, giving a final concentration of 10 µM Cu^2+^-TETAC. Co^2+^ did not appear to be compatible with TETAC, eliminating its cysteine reactivity. 1,4-Dithiothreitol (DTT) (Sigma-Aldrich, St. Louis, MO) was prepared as a 100 mM stock in water.

Absorption measurements were made using a Beckman Coulter DU 800 (Brea, CA). To measure the absorption of Cu^2+^ and Co^2+^ bound to HH, we used an α-helical peptide with the following sequence: Ac-ACAAKHAAKHAAAAKA-NH_2_, which was custom synthesized by Sigma-Aldrich (St. Louis, MO) (9). The peptide was dissolved in SBT to a final concentration of 1 mM. N-ethyl maleimide was prepared as a 200 mM stock in DMSO and added to the peptide solution at a final concentration of 2 mM to prevent the peptide from precipitating upon addition of Cu^2+^. To measure the absorption of Cu^2+^-TETAC, we used the unconjugated form (i.e. not bound to protein) at a concentration of 2 mM.

### Calculation of Quantum Yield

Because the emission spectrum of L-Anap is sensitive to its environment, we measured the emission spectrum and quantum yield for L-Anap incorporated into each of our two sites in MBP. The emission spectrum of MBP-322Anap was blue shifted relative to that of MBP-295Anap by about 20 nm, suggesting that the L-Anap in MBP-322 is in a somewhat more hydrophobic environment (*Fig. 13*A). To measure the quantum yield for each L-Anap position in MBP, we identified a solvent condition for free L-Anap that would mimic the environment of the fluorophores at each of the two sites. We measured the emission spectra of free L-Anap in SBT/ethanol mixtures that ranged from 0% to 100% ethanol. As shown in Fig. *Fig. 13*B, both the peak wavelength and amplitude of the emission spectra were sensitive to ethanol. We found that the emission spectrum of MBP-295TAG matched most closely to the emissions spectrum of free L-Anap in 20% ethanol (*Fig. 13*A, red) whereas the emission spectrum of MBP-322Anap matched most closely to free L-Anap in 85% ethanol (*Fig. 13*A, blue).

**Fig. 13.**
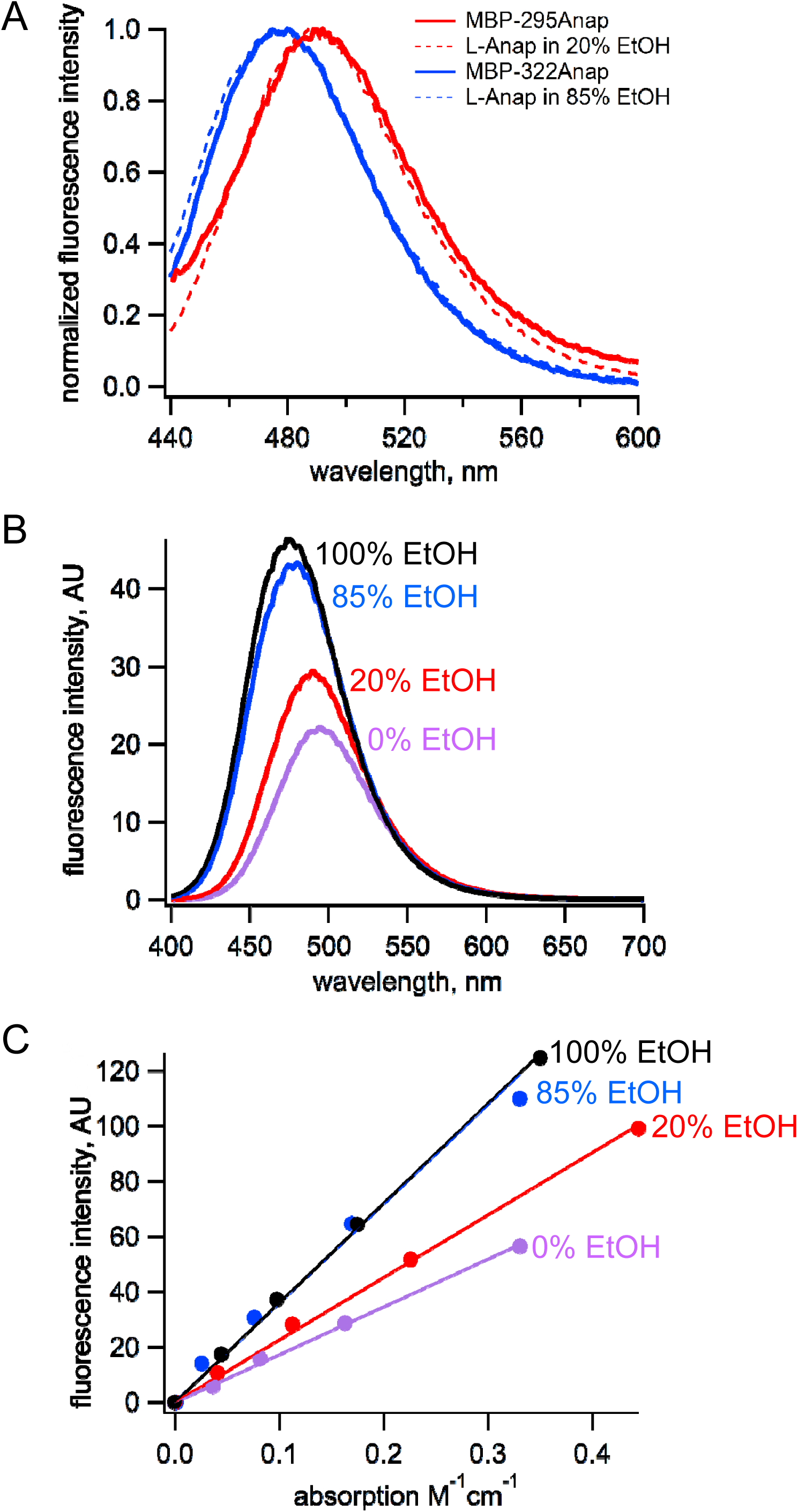
Estimation of quantum yield of Anap incorporate at the 295 and 322 sites in MBP. (A) Normalized emission spectra of MBP-295Anap (red line) and MBP-322Anap (blue line) showing the emission of Anap at 322 is blue shifted relative to that of Anap at 295. Also shown are the normalized emission spectra of free L-Anap in 20% ethanol (dashed red line), closely matching Anap at 295, and 85% ethanol (dashed blue line), closely matching Anap at 322. (B) Emission spectra of free L-Anap in different ethanol concentrations showing the environmental sensitivity of L-Anap. Over this range, as the environment becomes more hydrophobic, the peak fluorescence becomes larger and blue shifted. (C) Plot of fluorescence intensity versus absorption of free L-Anap in different ethanol concentrations. The quantum yield, the slow of these plots, increases with increasing ethanol concentration. The quantum yield of free L-Anap in 20% and 85% ethanol were used to estimate the quantum yield of Anap incorporate at the 295 and 322 sites, respectively.

Next, we measure the quantum yield of L-Anap in the different SBT/ethanol mixtures.

Samples of L-Anap were prepared in SBT, 20% ethanol in SBT, 85% ethanol in SBT, and 100% ethanol. The fluorescence intensity (excitation at 370 nm and emission at 480 nm) and absorption (at 370 nm) of each sample were measured as described above and plotted against each other in *Fig. 13*C. The quantum yield of L-ANAP in each of the mixtures was calculated relative to the reference quantum yield of L-ANAP in ethanol (0.48) (16) using the following equation (41):

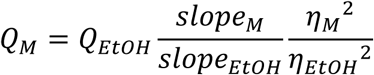

where *Q* is the quantum yield of L-ANAP, slope refers to the slope of the linear fits to the data, and *ɳ* is the refractive index. Quantum yield values were 0.23, 0.31, and 0.47 for 0%, 20%, and 85% ethanol, respectively. We therefore estimated the quantum yield for MBP-295Anap of 0.31 and for MBP-322Anap of 0.47. These estimates assume that EtOH:SBT mixtures mimic the L-Anap environment in MBP and that there are no endogenous quenchers within MBP.

### FRET Efficiency Analysis

For each time course experiment in the fluorometer, an averaged background trace from 6-8 experiments that did not contain protein, but to which Cu^2+^-TETAC and DTT, or Cu^2+^ and EDTA, were added, was subtracted from the protein-containing trace. The fraction of fluorescence quenching (F) was defined as follows:

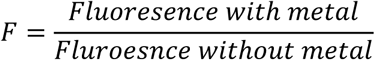

To determine the FRET efficiency, E, we corrected for other sources of energy transfer (e.g. solution quenching) using the following equations:

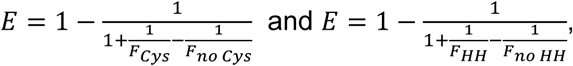

where F_Cys_ and F_no Cys_ are the fractional quenching by Cu^2+^-TETAC in protein with and without cysteines respectively, and F_HH_ and F_no HH_ are the fraction of quenching by Cu^2+^ in protein with and without HH sites, respectively. We also analyzed our data using simplified equations, which would account for nonspecific decreases in fluorescence (e.g. bleaching or loss of protein) but not energy transfer:

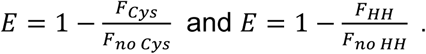

The values of E produced by the two analysis methods were similar because of the low degree of background quenching. For clarity, only the results of analysis with the former equations are shown. To calculate the mean and standard error of the mean for E, we used the mean and standard error of the mean for our F measurements (i.e. F_HH_, F_No HH_, F_Cys_, and F_No Cys_) in Monte Carlo resampling (1×10^6^ cycles; NIST Uncertainty Machine v1.3.4; (42)).

### Imaging and unroofing

Cells were imaged and unroofed as previously described (17, 20). Briefly, coverslips were mounted in a homemade chamber on a microscope stage and perfused for several minutes with HBR via gravity-flow perfusion. Poly-lysine solution (0.1 mg/mL 30,000-70,000 MW in PBS with 1 mM added CaCl_2_ and 1 mM added MgCl_2_) was then added to the chamber and the cells were incubated for 15 seconds. A 1:3 dilution of SBT was then added to the chamber, and the cells were incubated for 30 seconds. The solution was then replaced with SBT and the cells immediately exposed to a single pulse from a Branson Sonifier 450 with MicroTip with power = 2 and duty cycle = 50%. Unroofed cells were then perfused with SBT for at least two minutes to remove debris from the chamber before images were collected.

Experiments were performed using a Nikon Ti-E inverted microscope (Melville, NY) and a Nikon CFI Apo TIRF 60x oil immersion objective. Excitation light was provided by a Xenon arc lamp (model Lambda LS, Sutter Instrument Co., Novato, CA). For L-Anap excitation we used a 375/28 nm excitation filter and a 480/50 nm emission filter. Each experiment commenced with a pre-bleaching step, comprised of a seven-second exposure of the coverslip to L-Anap excitation light (20). Images for analysis were collected using exposures ranging from 100 ms to 500 ms, depending on sample intensity, using a QuantEM EMCCD camera (Photometrics, Tuscon, AZ) with readout speed of 5 MHz and the multiplier set to 20. Different metal concentrations and EDTA were applied via gravity perfusion. Cu^2+^-TETAC and DTT were applied directly to the chamber via a plastic Pasteur pipette. Five times the chamber volume was added in each case to ensure complete changeover of the solution.

### Image Analysis

Images were imported into ImageJ (43) for analysis. Regions of interest were drawn by hand to include most of a given cell, but excluding any regions that appeared only partially unroofed, which tended to occur at the edges. For each cell a background region was selected nearby that did not contain cells. The mean gray value of the background region of interest was subtracted from the mean gray value of the region of interest of the corresponding cell. This background subtraction was repeated for each image collected in an experiment using the same regions of interest. The mean and standard error of the mean for the FRET efficiency were calculated as described above.

### Distance calculations

The R_0_ values for each L-Anap site paired with each type of bound metal (i.e. Cu^2+^ or Cu^2+^-TETAC) were calculated using the measured emission spectrum of L-Anap at each site and the measured absorption spectra for each type of bound metal using the following equation (41):

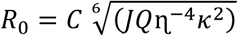

where *C* is a scaling factor, *J* is the normalized spectral overlap of the emission of the donor and absorption of the acceptor, *Q* is the quantum yield of L-Anap at the given site (see above), *ɳ* is the index of refraction (1.33 in our case), and κ^2^ is the orientation factor, assumed to be 2/3, a reasonable assumption for an anisotropic acceptor (15). Distances were calculated from the FRET measurements using the Fӧrster equation:

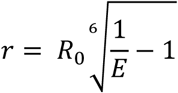

FRET efficiencies assuming a Gaussian distribution of distances between donor and acceptor, with FWHM = 8 Å (i.e. σ=3.4) were determined by numerically convolving the Fӧrster equation with the Gaussian function in Microsoft (Redmond, WA) Excel 2016. The corrected distances were then determined from plots of the FRET efficiency vs. the mean distance of the Gaussian distribution.

## Acknowledgements

Research reported in this publication was supported by the National Eye Institute of the National Institutes of Health under award numbers R01EY017564 (to SEG) and R01EY010329 (to WNZ), by the National Institute of Mental Health of the National Institutes of Health under award number R01MH102378 (to WNZ), by the National Institute of General Medical Sciences of the National Institutes of Health under award numbers R01GM100718 and R01GM125351 (to SEG and WNZ), and by the following additional awards from the National Institutes of Health: S10RR025429, P30DK017047, and P30EY001730. The authors declare no competing financial interests.

## Supplemental figure legends

**Supplemental Figure 1.**
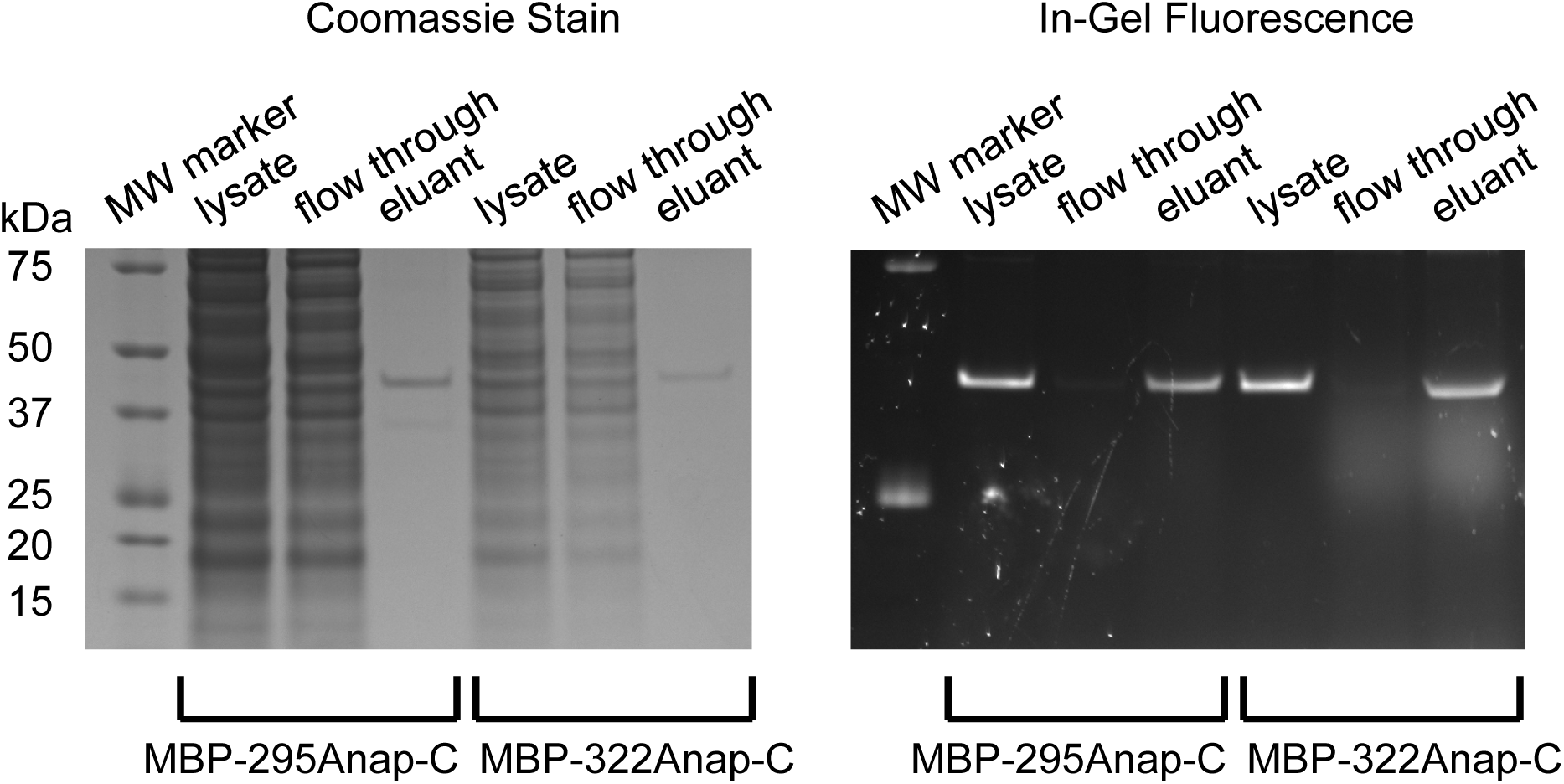
SDS-PAGE showing purification MBP constructs and specificity of Anap incorporation. Coomassie stained (left) and in-gel fluorescence (right) images of gel during the stages of purification of for MBP-295Anap and MBP-322Anap: cleared cell lysate, flow through, and elutant from M2-anti-FLAG beads. Similar fractions of the lysates were loaded for each stage, but 10 times more lysate was used for MBP-295Anap than MBP-322Anap. The absence of other bands in the lysate suggest strong specificity for Anap incorporation in MBP. Note, all constructs contain an N-terminal FLAG epitope, as described in Materials and Methods.

**Supplemental Figure 2.**
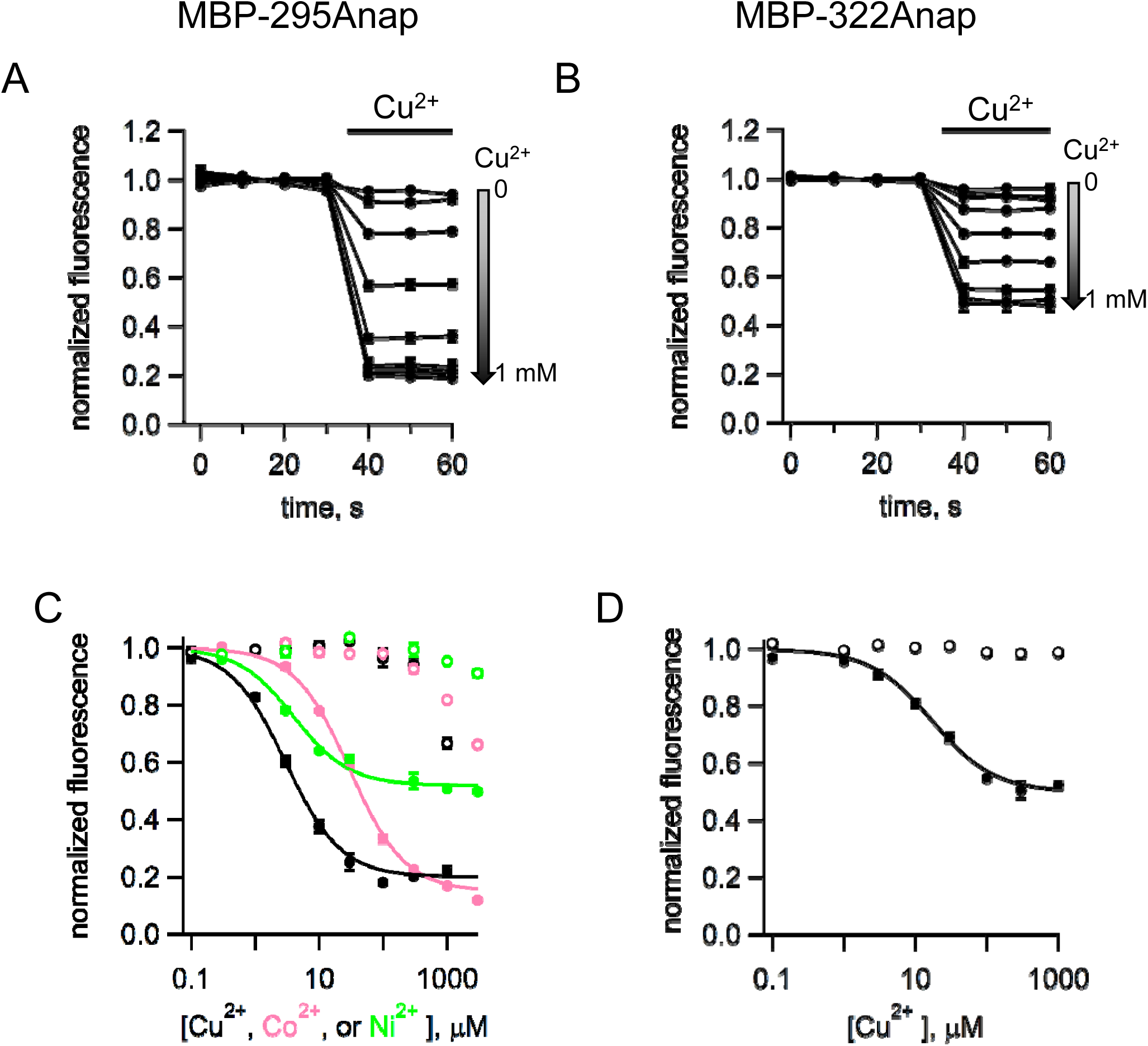
tmFRET with standard HH sites in MBP with different metals and metal concentrations. (A-B) Time course of tmFRET with HH sites for (A) MBP-295Anap-HH and (B) MBP-322Anap-HH with application of different concentrations of Cu^2+^ranging from 100 nM to 10 mM at the time of the bar. MBP-295Anap-HH was measured in the presence of 10 mM maltose and MBP-322Anap-HH was measure in the absence of maltose. The fluorescence of each construct was recorded every 10 s and normalized to the fluorescence in the absence of Cu^2+^. Shown are mean ± SEM for n = 3. (C-D) Plots of the concentration dependence of metal quenching for (C) MBP-295Anap and (D) MBP-322Anap. The quenching in shown for Cu^2+^ (black), Ni^2+^ (green), and Co^2+^ (magenta) for constructs with (filled symbols) or without (open symbols) a HH site. Shown are mean ± SEM for n = 3. The fits to the data correspond to a single binding isotherm and indicate that MBP-295Anap-HH has an affinity of 3 µM, 4 µM, and 30 µM for Cu^2+^, Ni^2+^, and Co^2+^ respectively, and MBP-322Anap-HH has an affinity of 17 µM for Cu^2^. The fractional fluorescence at saturating concentration for MBP-295Anap-HH was 0.2, 0.48, and 0.15 for Cu^2+^, Ni^2+^, and Co^2+^ respectively, and for MBP-322Anap-HH was 0.5 for Cu^2^.

Supplemental Movie 1 – Conformation change in MBP upon maltose binding. Morph of the X-ray crystal structures of MBP in the apo (PDB ID: 1N3W) and maltose-bound (PDB ID: 1N3X) states. Shown are the measurements of the distances for the FRET pairs used in this study, MBP-295Anap-C (top) and MBP-322Anap-C (bottom). Maltose is shown in magenta.

## References

1. Henzler-Wildman K & Kern D (2007) Dynamic personalities of proteins. Nature 450(7172):964–972.

2. Cheng Y (2015) Single-Particle Cryo-EM at Crystallographic Resolution. Cell 161(3):450–457.

3. Taraska JW & Zagotta WN (2010) Fluorescence applications in molecular neurobiology. Neuron 66(2):170–189.

4. Stryer L & Haugland RP (1967) Energy transfer: a spectroscopic ruler. Proc Natl Acad Sci U S A 58(2):719–726.

5. Best RB, et al. (2007) Effect of flexibility and cis residues in single-molecule FRET studies of polyproline. Proc Natl Acad Sci U S A 104(48):18964–18969.

6. Schuler B, Lipman EA, Steinbach PJ, Kumke M, & Eaton WA (2005) Polyproline and the “spectroscopic ruler” revisited with single-molecule fluorescence. Proc Natl Acad Sci U S A 102(8):2754–2759.

7. Yu X, Wu X, Bermejo GA, Brooks BR, & Taraska JW (2013) Accurate high-throughput structure mapping and prediction with transition metal ion FRET. Structure 21(1):9–19.

8. Taraska JW, Puljung MC, Olivier NB, Flynn GE, & Zagotta WN (2009) Mapping the structure and conformational movements of proteins with transition metal ion FRET. Nat Methods 6(7):532–537.

9. Taraska JW, Puljung MC, & Zagotta WN (2009) Short-distance probes for protein backbone structure based on energy transfer between bimane and transition metal ions. Proc Natl Acad Sci U S A 106(38):16227–16232.

10. Horrocks WD, Holmquist B, & Vallee BL (1975) Energy transfer between terbium (III) and cobalt (II) in thermolysin: a new class of metal--metal distance probes. Proc Natl Acad Sci U S A 72(12):4764–4768.

11. Latt SA, Auld DS, & Vallee BL (1972) Distance measurements at the active site of carboxypeptidase A during catalysis. Biochemistry 11(16):3015–3022.

12. Richmond TA, Takahashi TT, Shimkhada R, & Bernsdorf J (2000) Engineered metal binding sites on green fluorescence protein. Biochem Biophys Res Commun 268(2):462–465.

13. Sandtner W, Bezanilla F, & Correa AM (2007) In vivo measurement of intramolecular distances using genetically encoded reporters. Biophys J 93(9):L45–47.

14. Latt SA, Auld DS, & Valee BL (1970) Surveyor substrates: energy-transfer gauges of active center topography during catalysis. Proc Natl Acad Sci U S A 67(3):1383–1389.

15. Selvin PR (2002) Principles and biophysical applications of lanthanide-based probes. Annu Rev Biophys Biomol Struct 31:275–302.

16. Chatterjee A, Guo J, Lee HS, & Schultz PG (2013) A genetically encoded fluorescent probe in mammalian cells. J Am Chem Soc 135(34):12540–12543.

17. Zagotta WN, Gordon MT, Senning EN, Munari M, & Gordon SE (2016) Measuring distances between TRPV1 and the plasma membrane using a noncanonical amino acid and transition metal ion FRET. Journal of General Physiology 147(2):201–216.

18. Sobakinskaya E, Schmidt Am Busch M, & Renger T (2018) Theory of FRET “Spectroscopic Ruler” for Short Distances: Application to Polyproline. J Phys Chem B 122(1):54–67.

19. Heuser J (2000) The production of ‘cell cortices′ for light and electron microscopy. Traffic 1(7):545–552.

20. Gordon SE, Senning EN, Aman TK, & Zagotta WN (2016) Transition metal ion FRET to measure short range distances at the intracellular surface of the plasma membrane. Journal of General Physiology 147(2):189–200.

21. Kalstrup T & Blunck R (2013) Dynamics of internal pore opening in K(V) channels probed by a fluorescent unnatural amino acid. Proc Natl Acad Sci U S A 110(20):8272–8277.

22. Aman TK, Gordon SE, & Zagotta WN (2016) Regulation of CNGA1 channel gating by interactions with the membrane. Journal of Biological Chemistry.

23. Schmied WH, Elsässer SJ, Uttamapinant C, & Chin JW (2014) Efficient multisite unnatural amino acid incorporation in mammalian cells via optimized pyrrolysyl tRNA synthetase/tRNA expression and engineered eRF1. J Am Chem Soc 136(44):15577–15583.

24. Dai G, James ZM, & Zagotta WN (2018) Dynamic rearrangement of the intrinsic ligand regulates KCNH potassium channels. J Gen Physiol.

25. Dai G & Zagotta WN (2017) Molecular mechanism of voltage-dependent potentiation of KCNH potassium channels. Elife 6.

26. Puljung MC & Zagotta WN (2013) A secondary structural transition in the C-helix promotes gating of cyclic nucleotide-regulated ion channels. J Biol Chem 288(18):12944–12956.

27. Arnold FH & Haymore BL (1991) Engineered metal-binding proteins: purification to protein folding. Science 252(5014):1796–1797.

28. Suh SS, Haymore BL, & Arnold FH (1991) Characterization of His-X3-His sites in alpha-helices of synthetic metal-binding bovine somatotropin. Protein Eng 4(3):301–305.

29. Martineau P, Szmelcman S, Spurlino JC, Quiocho FA, & Hofnung M (1990) Genetic approach to the role of tryptophan residues in the activities and fluorescence of a bacterial periplasmic maltose-binding protein. J Mol Biol 214(1):337–352.

30. Farnsworth CC, Wolda SL, Gelb MH, & Glomset JA (1989) Human lamin B contains a farnesylated cysteine residue. J Biol Chem 264(34):20422–20429.

31. Anderegg RJ, Betz R, Carr SA, Crabb JW, & Duntze W (1988) Structure of Saccharomyces cerevisiae mating hormone a-factor. Identification of S-farnesyl cysteine as a structural component. J Biol Chem 263(34):18236–18240.

32. Jeschke G (2012) DEER distance measurements on proteins. Annu Rev Phys Chem63:419–446.

33. Frauenfelder H, Sligar SG, & Wolynes PG (1991) The energy landscapes and motions of proteins. Science 254(5038):1598–1603.

34. Sakata S, Jinno Y, Kawanabe A, & Okamura Y (2016) Voltage-dependent motion of the catalytic region of voltage-sensing phosphatase monitored by a fluorescent amino acid. Proc Natl Acad Sci U S A 113(27):7521–7526.

35. Wulf M & Pless SA (2018) High-Sensitivity Fluorometry to Resolve Ion Channel Conformational Dynamics. Cell Rep 22(6):1615–1626.

36. Soh MS, Estrada-Mondragon A, Durisic N, Keramidas A, & Lynch JW (2017) Probing the Structural Mechanism of Partial Agonism in Glycine Receptors Using the Fluorescent Artificial Amino Acid, ANAP. ACS Chem Biol 12(3):805–813.

37. Wen M, et al. (2015) Site-specific fluorescence spectrum detection and characterization of hASIC1a channels upon toxin mambalgin-1 binding in live mammalian cells. Chem Commun (Camb) 51(38):8153–8156.

38. Cunningham TF, et al. (2015) Cysteine-specific Cu2+ chelating tags used as paramagnetic probes in double electron electron resonance. J Phys Chem B 119(7):2839–2843.

39. Sengupta I, et al. (2015) Protein structural studies by paramagnetic solid-state NMR spectroscopy aided by a compact cyclen-type Cu(II) binding tag. J Biomol NMR 61(1):1–6.

40. Bjerrum OJ-N, C (1986) Buffer systems and transfer parameters for semidry electroblotting with a horizontal apparatus. Electrophoresis ‘86, ed Dunn M (VCH, London), pp 315–327.

41. Lakowicz JR (2006) Principles of fluorescence spectroscopy (Springer, New York) 3rd Ed pp xxvi, 954 pages.

42. Lafarge T & Possolo A (2016) The NIST Uncertainty Machine. The Journal of Measurement Science 10(3):20–27.

43. Schneider CA, Rasband WS, & Eliceiri KW (2012) NIH Image to ImageJ: 25 years of image analysis. Nat Methods 9(7):671–675.

